# DUSP6 mediates resistance to JAK2 inhibition and drives leukemic progression

**DOI:** 10.1101/2021.06.27.449656

**Authors:** Tim Kong, Angelo Brunelli Albertoni Laranjeira, Kangning Yang, Daniel A. C. Fisher, LaYow Yu, Anthony Z Wang, Marianna Ruzinova, Jared S. Fowles, Maggie J. Allen, Hamza Celik, Grant A. Challen, Sidong Huang, Stephen T. Oh

**Author notes:** These authors contributed equally (Angelo Brunelli Albertoni Laranjeira, Kangning Yang).

## Abstract

Chronic myeloproliferative neoplasms (MPNs) exhibit a propensity for transformation to secondary acute myeloid leukemia (sAML), for which the underlying mechanisms remain poorly understood, resulting in limited treatment options and dismal clinical outcomes. Here, we performed bulk transcriptome profiling accompanied by single cell RNA-sequencing on CD34+ stem/progenitor cells from serial patient samples obtained at the chronic MPN and sAML phases, and identified aberrantly increased expression of dual-specificity phosphatase 6 (*DUSP6*) underlying disease transformation. Genetic and pharmacologic targeting of DUSP6 led to inhibition of S6 and JAK/STAT signaling, resulting in potent suppression of cell proliferation, while also reducing inflammatory cytokine production in primary samples. Furthermore, ectopic *DUSP6* expression augmented proliferation and mediated JAK2 inhibitor resistance, while DUSP6 inhibition reduced colony-forming potential of JAK2 inhibitor-persistent patient cells. Mechanistically, DUSP6 perturbation dampened S6 signaling via inhibition of RSK1, which we identified as a second indispensable candidate associated with poor clinical outcome. Lastly, DUSP6 inhibition potently suppressed disease development across *Jak2* V617F and *MPL* W515L MPN mouse models, and sAML patient-derived xenografts. These findings underscore DUSP6 in driving disease transformation and therapeutic resistance, and highlight the DUSP6-RSK1 axis as a novel, druggable pathway in myeloid malignancies.

## Introduction

Myeloproliferative neoplasms (MPNs) are clonally derived from hematopoietic stem/progenitor cells (HSPCs) and typically harbor somatic mutations in one of three genes (*JAK2*, *CALR*, *MPL*) leading to aberrant activation of JAK-STAT signaling(1–3). Polycythemia vera (PV) and essential thrombocythemia (ET) are chronic MPNs characterized by overproduction of blood cells (erythrocytosis, thrombocytosis) and thrombohemorrhagic complications, while myelofibrosis (MF) is a more aggressive MPN subtype characterized by anemia, splenomegaly, bone marrow fibrosis, and constitutional symptoms (fever, night sweats, fatigue, weight loss)(4). Small molecule inhibitors of JAK2 provide symptomatic benefit for MPN patients, but they do not eradicate the underlying malignant clone, nor do they reliably prevent disease progression(5, 6). These findings suggest that other signaling pathways may be dysregulated, promoting survival of the malignant clone despite JAK2 inhibition.

Chronic MPNs exhibit a propensity for transformation to secondary acute myeloid leukemia (sAML), which imparts a dismal prognosis with a median survival of < 6 months(7). Leukemic transformation is often associated with the acquisition of additional mutations (e.g. *TP53*), and several studies have established that the presence of “high-risk” mutations (e.g. *ASXL1*, *EZH2*, *SRSF2*, *IDH1/2*) at the chronic MPN stage confers an increased risk of sAML transformation(7–10). However, the manner in which these genetic alterations contribute to leukemic transformation remains poorly understood. Despite vigorous research, therapies capable of reliably halting or preventing MPN disease progression remain elusive. Therefore, there is a pressing need to better define the mechanisms responsible for transformation of MPNs to sAML, which may translate to therapies capable of preventing and/or treating disease progression.

There has been emerging interest in the role and targeting of the MAPK/ERK signaling pathway in hematological malignancies(11, 12). DUSP6 is a member of the MAPK phosphatase family that is categorized into three subgroups based on sequence homology, subcellular localization, and substrate specificity(13–15). DUSP6 and other Class II DUSPs are highly selective for ERK while class I DUSPs (including DUSP1) act on JNK and p38 (in addition to ERK). Canonically, DUSP6 regulates signaling through ERK (and downstream signaling mediators) via a dynamic feedback regulatory axis: activation of MAPK signaling leads to induction of DUSP6 expression, followed by interaction of DUSP6 with ERK leading to dephosphorylation and inactivation of ERK(15). The specific role of DUSP6 is largely context dependent, such that aberrant DUSP6 expression may exert seemingly paradoxical effects on ERK and related signaling pathways, as illustrated by prior studies supporting an oncogenic role for DUSP6 in some cancers, but a tumor suppressor role in other settings(14, 15). Given these observations, DUSP6 may be an important mediator of disease burden and therapeutic response.

Here, using multiomic approaches, we identify DUSP6 to be a critical driver of MPN disease progression, whose aberrant expression also confers resistance to JAK2 inhibition. We further exploit this dependency pharmacologically via DUSP6 targeting across *in vitro*, *in vivo,* and *ex vivo* MPN/sAML models.

## Methods

### Cell culture

HEL (ATCC) and UKE-1 (Coriell Institute) cells were cultured in RPMI 1640 Medium ATCC modification and RPMI 1640 Medium (ThermoFisher, Waltham, MA), respectively, supplemented with 10% fetal bovine serum (FBS) and 1% penicillin/streptomycin. All cell lines were maintained at 37°C and 5% CO2 and regularly tested for mycoplasma.

### Patient samples

Patient and healthy donor control peripheral blood (PB) or bone marrow (BM) samples were obtained with written consent according to a protocol approved by the Washington University Human Studies Committee (WU no. 01-1014). Mononuclear cells (PBMC or BMMC) were obtained by Ficoll gradient extraction and cryopreserved according to standard procedures. Additional BMMCs were purchased from AllCells (Alameda, California). A list of patient samples utilized in this study is provided in Table 1.

### Compounds and antibodies

Ruxolitinib, fedratinib, and trametinib were purchased from Selleck Chemicals (Houston, TX). BCI and BI-D1870 were purchased from MedChemExpress (Monmouth Junction, NJ). For *in vivo* treatment, BCI-HCl and ruxolitinib were purchased from MedChemExpress. Human TPO was purchased from Peprotech (Rocky Hill, NJ). Protein Transport Inhibitor Cocktail (500X) was purchased from ThermoFisher.

Immunobloting antibodies were used as follows: DUSP6 (ab76310) from Abcam (Cambridge, MA). pRB S795 (9301), cleaved caspase-3 Asp175 (9661), pERK1/2 T202/7204 (9101), RSK1 (8408), p90RSK T359/S363 (9344), S6 (2317), pS6 S235/236 (4858), pSTAT3 Y705 (9131), pSTAT3 S727 (6E4), and pSTAT5 Y694 (9351) from Cell Signaling Technology (Danvers, MA). HSP90 (sc-13119), ERK1 (sc-93), and ERK2 (sc-154) from Santa Cruz Biotechnology (Dallas, TX). KLF2 (SAB1403063) from Sigma-Aldrich (St. Louis, MO).

Flow cytometry and cell sorting antibodies were used as follows and were from BD Biosciences (Franklin Lakes, NJ) : Alexa Fluor 488 Mouse Anti-Stat3 (pY705), PE-Cy7 Mouse anti-Stat5 (pY694), FITC Mouse Anti-Human CD34 (8G12), APC Mouse Anti-Human CD38 (HIT2), PE Mouse Anti-Human CD3 (UCHT1), PE Mouse Anti-Human CD19 (HIB19), PE Mouse Anti-Human CD14 (M5E2), PE Mouse Anti-Human CD16 (3G8), and PE Mouse Anti-Human CD7 (M-T701).

### Lentiviral transduction and constructs

Lentiviral transduction with low multiplicity of infection (MOI) was performed using the protocol as described at http://www.broadinstitute.org/rnai/public/resources/protocols. Transduced cells were selected in puromycin or blasticidin for 2-4 days and plated for downstream assays immediately after selection.

Individual shRNA and ORF vectors used were from the Mission TRC library (Sigma), and ORF collections were developed by members of the ORFeome Collaboration (Sigma/TransOMIC), provided by the McGill Platform for Cellular Perturbation (MPCP) of Rosalind and Morris Goodman Cancer Research Centre at McGill University: control vector pLKO.5, sh*DUSP6* #1 (TRCN0000233473), sh*DUSP6* #2 (TRCN0000233471), sh*RPS6KA1* #1 (TRCN0000001386), sh*RPS6KA1* #2 (TRCN0000001388), sh*KLF2* #1 (TRCN0000020727), sh*KLF2* #2 (TRCN0000020728), control vector pLX304-e*GFP*, pLX304-*DUSP6* (ccsbBroad304_00468), and pLX304-*RPS6KA1* (ccsbBroad304_06896).

### Cell viability assays

HEL, UKE-1, and patient PBMCs were seeded into 96-well plates at a density of 2 × 10^3^ to 2 × 10^4^ cells/well upon gene perturbation or treated with indicated doses of inhibitors and cultured for 72-96 hours. At endpoint, AlamarBlue viability reagent was added and absorbance was read in a BioTek microplate reader.

### Generation of persistent HEL cells

HEL cells were cultured in the presence of 1uM of ruxolitinib or fedratinib for two months. Cells were pelleted and resuspended in 1uM of inhibitor weekly to maintain selection pressure.

### Immunoblotting

5 × 10^5^ to 2 × 10^6^ cells/well were seeded into 6-well or 12-well plates. After the indicated treatment duration, cells were washed with ice-cold PBS and lysed with protein sample buffer. Samples were processed with NuPAGE Novex Gel Electrophoresis Systems (ThermoFisher) followed by standard immunoblotting procedure.

### qRT-PCR

RNA from plated cells were extracted with TRIzol (ThermoFisher). cDNA was synthesized using the Maxima First-Strand cDNA Synthesis Kit (ThermoFisher). Relative mRNA levels were measured through qRT-PCR using SYBR Green Master Mix (Roche) and normalized to the expression of β-actin (*ACTB*). Primer sequences are as follows:

*ACTB* forward, 5′-GTTGTCGACGACGAGCG-3′

*ACTB* reverse, 5′-GCACAGAGCCTCGCCTT-3′

*DUSP6* forward, 5’-GTAGGGGTTGTGAATTGTGT-3’

*DUSP6* reverse, 5’-ACCACCAATACCCACAACCA-3’

### Flow cytometry, annexin V apoptosis assay, and cell cycle assay

#### Signaling flow cytometry

2.5 x 10^5^ HEL cells were seeded and treated with 1 µM BCI for 24 hours. Post treatment, cells were fixed with paraformaldehyde, washed with PBS, permeabilized with methanol, and stained with flow antibodies. Samples were run on BD FACSCanto II Cell Analyzer utilizing unstained controls, and downstream analysis was performed with FlowJo (Ashland, OR). Cell gating is as follows: SSC-A/FSC-A (size, live/dead), FITC-A/PE-Cy7-A (pSTAT3 Y705/pSTAT5 Y694).

#### Annexin V apoptosis assay

HEL cells were seeded at 4 x 10^5^ cells/well and BCI, BI-D1870, or DMSO control were added for their indicated treatment duration. Protocol was followed per the PE Annexin V Apoptosis Detection Kit I (BD Biosciences). In brief, post treatment, cells were washed with PBS and resuspended in binding buffer, incubated with PE-Annexin V, and stained with 7-AAD viability staining solution. Cells were then analyzed by flow cytometry, in which AnnexinV+/7-AAD+ cells were identified as necroptotic, AnnexinV+/7-AAD- as apoptotic, and AnnexinV-/7-AAD- as healthy.

#### Cell cycle assay

HEL cells were seeded at 4 x 10^5^ cells/well and BCI, BI-D1870, or control were added for the indicated duration. Post treatment, cells were washed with PBS and fixed in 70% ethanol for 24 hours. Cells were washed with PBS and then treated with RNase, followed by staining with propidium iodide. Cells were then analyzed by flow cytometry.

### CD34+ cell patient microarray

CD34+ cells from MF (n= 14) and sAML (n = 6) patient PBMCs and from normal BMMCs (n = 5) were sorted. Cryopreserved cells were thawed, filtered, and stained with the following immunostaining panel for sorting: CD34, CD38, CD3, CD19, CD14, and CD16 antibodies, in addition to 7-AAD viability marker. 7AAD- lin- CD34+ cells were sorted and mRNA was isolated, amplified, and converted to biotinylated sense-strand cDNA with WT Pico Reagent Kit (ThermoFisher). cDNA were then probed by GeneChipHuman Gene 2.0 ST Arrays (Affymatrix). Raw files were processed with the oligo package, quality controlled with AUC and RLE cut-offs, and normalized using the RMA-normalized log2-transformed algorithm. Differential gene expression analysis comparing sAML vs MF CD34+ samples was performed with Limma(16), where a Benjamini-Hochberg adjusted p-value cut-off of 0.05 was used.

### HEL RNA sequencing

RNA from HEL parental, HEL-RuxR, Hel Fed-R, HEL 4h control, HEL 24h control, HEL 4h BCI, HEL 24h BCI, a second set of HEL 24h control, HEL 24h BI-D1870, and HEL 24h BCI + BI-D1870 were extracted using RNeasy Mini Kit (Qiagen) with DNase treatment in duplicate. Samples were then indexed, pooled, and sequenced by Illumina NovaSeq 6000. Basecalls and demultiplexing were performed with Illumina’s bcl2fastq software and a custom python demultiplexing program with a maximum of one mismatch in the indexing read. RNA-seq reads were then aligned to the Ensembl release 76 primary assembly with STAR version 2.5.1a(17). TMM normalization size factors in EdgeR(18) were calculated to adjust for samples for differences in library size. Ribosomal genes and genes not expressed in the smallest group size minus one samples greater than one count-per-million were excluded from further analysis. The TMM size factors and the matrix of counts were then imported into the R/Bioconductor package Limma(16). Differential expression analysis was then performed to analyze for differences between conditions and the results were filtered for only those genes with Benjamini-Hochberg false-discovery rate adjusted p-values less than or equal to 0.05. Gene set enrichment analysis (GSEA) of MSigDb pathways was performed using the R/Bioconductor package GAGE(19) to test for changes in expression of the reported log 2 fold-changes reported by Limma in each term versus the background log 2 fold-changes of all genes found outside the respective term, and using GSEA(20) software with the Hallmark gene set, KEGG signaling and metabolism, or other curated gene sets to identify altered pathways.

### Single cell RNA-seq with surface TotalSeqA protein expression detection

Cryopreserved BMMCs from two normal control donors (N34 and N39), and PBMCs from serial collections from three patients, including chronic MPN phase and subsequent sAML phase were utilized: Patient 381812 (MF post ET and sAML 12 months following), Patient 145790 (two collections from MF post ET separated by 30 months and sAML 25 months following), and Patient 374024 (chronic PV and sAML 6 months following). Non-lymphoid CD34 cells were identified by the following immunostaining panel: CD34, CD38, CD3, CD7, CD19, and 7AAD. 7AAD- CD3- CD7- CD19- CD34+ cells were sorted and subsequently labeled with a panel of oligonucleotide conjugated antibodies (Table 9) according to TotalSeqA protocol from 10X Genomics (Pleasanton, CA). Oligonucleotide labeled cells were subjected to droplet bead capture, followed by cell lysis, reverse transcription, and amplification following protocol. Single cell derived cDNA libraries were sequenced on NovaSeq S4 cell sequencer (Illumina, San Diego, CA). Cellranger (10x Genomics, version 3.0.1) was used for transcript alignment, counting, and inter-library normalization of single cell samples. Seurat(21) (version 4.0.0) was used for cluster analysis. Cells containing fewer than 500 genes, a nCount value greater than the nCount value of the 93rd percentile of the entire sample, and more than 10% mitochondrial transcripts were removed, after which all filtered samples were merged. RNA reads were normalized with LogNormalize, ADT antibodies for surface protein expression were normalized with Centered Log-Ratio method, and data were scaled. Cell cycle phase was determined using CellCycleScoring, based upon on relative expression of phase-specific genes(22). Principle component analysis was performed on variable genes and the optimal number of PCs was selected to be 44 after JackStrawPlot analysis. Dimensionality reduction and visualization were performed with the uniform manifold approximation and projection (UMAP) algorithm and unsupervised graph-based clustering of cells was performed at a resolution of 0.8. Differential gene expression analysis between populations/samples of interest was performed utilizing FindAllMarkers with Wilcoxon rank sum test. Gene set enrichment analysis of DEGs was run using fgsea(23) and the Hallmark gene set from msigdbr. Subsequent subsetting of samples was performed for downstream analyses as indicated. Trajectory analysis was performed with Monocle 2(24) (version 2.18.0) utilizing DEGs from each cell cluster. Dimensionality reduction was performed with reduceDimension method DDRTree and custom root state was set to HSC or earliest pluripotent progenitor cell population. DEG analysis was utilized to identify branch-specific genes at each branchpoint post-trajectory analysis. Pseudotime expression of genes of interest was performed across chronic MPN and sAML samples, and along disease states from trajectory analysis. Cell populations were identified using ADT antibody markers and gene set signatures provided in Supplementary Fig S4.

### CD34+ cell colony assay

MPN and sAML patient PBMCs and normal BMMCs samples were thawed, filtered, and stained with CD34, CD38, CD3, CD19, CD14, and CD16 antibodies, in addition to 7-AAD viability marker. 7AAD- lin- CD34+ cells were sorted and seeded at 1,000 cells/mL in Methocult H4034 (StemCell) with the indicated inhibitors. Each condition was plated in duplicate or triplicate as indicated. Myeloid and erythroid colonies were counted after 10-14 days post-seeding.

### Mass cytometry (CyTOF)

Signaling and cytokine CyTOF experiments utilizing HEL cells and patient PBMCs and BMMCs were conducted on a CyTOF2 mass cytometry (Fluidigm) with validated antibody panels and following protocols as previously described(25–29). In brief, primary cells were thawed, filtered, and counted on a hemocytometer. For primary sample signaling CyTOF, cells were then labelled with cisplatin, washed, and incubated with indicated stimulant or inhibitor for the indicated duration. Cells were then fixed with paraformaldehyde, permeabilized with perm buffer, and barcoded. Following, cells were stained with surface marker antibodies, permeabilized with methanol, stained with intracellular antibodies, and resuspended in DNA IR-intercalator. For primary cytokine CyTOF, cells were incubated with BCI and TPO for 4 hours, and protein transport inhibitor cocktail (eBioscience) was added at the 2 hour treatment mark. Following, cells were stained with surface markers antibodies, stained with cisplatin, and fixed with paraformaldehyde. Cells were then permeabilized with eBioscience perm buffer, stained with intracellular antibodies, fixed, barcoded, and suspended in Ir-intercalator. For persistent HEL cell signaling CyTOF, cells were labeled with cisplatin, treated with indicated inhibitors for 1 hour, fixed, barcoded, and permeabilized with methanol. Following, cells were stained with intracellular antibodies resuspended in Ir-intercalator. For sAML14 PDX CyTOF, the femurs of transplanted NSGS mice at end point were harvested and flushed, and bone marrow cells were collected and filtered. Cells were then labelled with cisplatin, washed, fixed, permeabilized, and barcoded. Following, cells were stained with surface marker antibodies, permeabilized with methanol, stained with intracellular antibodies, and resuspended in DNA IR-intercalator. Post CyTOF run, cell identities were debarcoded and data was analyzed in Cytobank (cytobank.org) and FlowJo (TreeStar, Eugene, OR). Lin- negative CD34+ cells were gated as follows: CD3-, CD19-, CD235-, CD71-, CD61-, CD14-, CD15-, CD16-, CD34+. For sAML PDX14, to separate human and mouse bone marrow cells, post CyTOF run cells were gated by surface expression of hCD45 and mCD45. CyTOF antibody panels are provided in Table 9.

### Immunofluorescence and immunohistochemistry

Formalin fixed paraffin embedded (FFPE) blocks prepared from patient bone marrow core biopsies were sectioned to 5µm thickness on slides. Slides were deparaffinized by heating to 60°C for 2 hours, followed by serial exposure to xylene, 100%, 95%, 80%, and 70% ethanol solutions, and water. Rehydrated slides were treated with 10mM Tris, 1mM EDTA, pH 9.0 for 30 minutes at 96°C for antigen retrieval. Sections were blocked with 3% bovine serum albumin in phosphate buffered saline (PBS) for 30 minutes at room temperature. Sections were stained with mouse antibody to human DUSP6 antibody for 16-20h at 4°C, followed by washes with 0.5% Triton X-100 in PBS, PBS alone, and labeling with anti-mouse secondary antibody, Alexa 647 conjugate for 1h at room temperature. Sections were counterstained with DAPI to visualize cell nuclei, and imaged on a Leica DM6 upright fluorescent microscope, using a Leica DFC 700T camera and LAS-X imaging suite.

For mouse histopathology, tibias were extracted and fixed in 10% neutral buffered formalin overnight at 4°C. Post fixation, tibias were decalcified in 14% EDTA for 12 days and then rehydrated with serial exposure to 20%, 30%, 50%, and 70% ethanol, all for 1 hour. Tibias were then rinsed in PBS and processed for paraffin embedding and sectioned at 5 μM. H&E staining was performed by the Washington University Musculoskeletal Histology and Morphometry Core. Reticulin staining of bone marrow samples was performed on Ventana BenchMark Special Stains automated slide system (Ventana Medical Systems, Tucson, AZ, USA).

### Imaging mass cytometry

Cryopreserved PBMCs from healthy control LRS2, MF20, and sAML1 were pelleted, fixed in 1.6% formaldehyde for 10 minutes at room temperature followed by 10% neutral buffered formalin for 48h at 4°C, and embedded in Histogel (ThermoFisher), followed by subsequent embedding in paraffin and sectioning, rehydration, antigen retrieval, and blocking as described for immunofluorescence. Antibodies for imaging mass cytometry were conjugated to metal-chelator complexes according to an established conjugation kit protocol (Fluidigm). Mass cytometry signals were recorded with the Hyperion imaging mass cytometry platform (Fluidigm). Exported mcd files were loaded into histoCAT++(30) and a median filter (size 3) was applied to all channels before image export. Antibodies panels are provided in Table 9.

### *In vivo* mouse models

All procedures were conducted in accordance with the Institutional Animal Care and Use Committee of Washington University.

#### Normal mice

PBS vehicle or BCI-HCl (25 mg/kg) was administered intraperitonially once a day to 7 week-old C57BL/6 mice. Mice were treated for four weeks and peripheral blood was collected every Friday and cells were analyzed by Hemavet. Mice were sacrificed at endpoint, and body, spleen, and liver weights were recorded.

#### *Jak2* V617F model

Whole bone marrow was harvested from femurs and tibias of CD45.2 donor mice (6-12 weeks old) by flushing. Cells were resuspended and filtered in sort buffer (PBS containing 0.5% BSA and 2mM EDTA). 1 x 10^5^ bead-enriched Kit+ cells were collected from CD45.2 *Jak2* V617F donor mice and transplanted via tail vein into CD45.1 lethally irradiated recipient mice. Recipient mice were irradiated with a total of 1,100 cGy, given as two separate doses of 550 cGy with 3-5 hours between doses. Within three hours of the last irradiation, mice were transplanted. For two weeks following transplant, the mice were caged with water containing 0.5 mg/mL Trimethoprim/sulfamethoxazole (Washington University Division of Comparative Medicine Pharmacy). Two weeks after transplant, peripheral blood was collected, and hematologic parameters were measured by Hemavet and flow cytometry. Mice were treated with PBS vehicle or BCI-HCl (25 mg/kg) intraperitonially daily. Treatment began two weeks after transplant, Monday through Friday. Mice were treated for four weeks and peripheral blood was collected every Friday and cells were analyzed by Hemavet and flow cytometry. Mice were sacrificed at endpoint, and body, spleen, and liver weights were recorded.

#### *MPL* W515L model

Retroviral transduction was performed as described previously(31). Briefly, retrovirus was produced by transfecting 293T cells with viral plasmids using jetPrime transfection reagent and viral supernatant was collected after 24 hours. Whole bone marrow was harvested from femurs and tibias of CD45.2 donor mice (6-12 weeks old) by flushing. The cells were resuspended and filtered in sort buffer (PBS containing 0.5% BSA and 2mM EDTA). Isolated bead-enriched Kit+ cells were grown in 6-well plates for 2 days while undergoing spinoculation with the viral supernatant once per day by centrifugation at 350 x g for 30 minutes. GFP expression was verified by flow cytometry prior to transplantation. 5 x 10^5^ bead-enriched Kit+ cells retrovirally transduced with *MPL* W515L were transplanted via tail vein into CD45.1 lethally irradiated recipient mice (7 weeks old). Recipient mice were irradiated with a total of 1,100 cGy, given as two separate doses of 550 cGy with 3-5 hours between doses. Within three hours of the last irradiation, mice were transplanted. For two weeks following transplant, the mice were caged with water containing 0.5 mg/mL Trimethoprim/sulfamethoxazole. Two weeks after transplant, peripheral blood was collected and hematologic parameters were measured by Hemavet and flow cytometry. Mice were treated with PBS vehicle or BCI-HCl (25 mg/kg) intraperitonially daily. Treatment began two weeks after transplant, Monday through Friday. Mice were treated for three weeks and peripheral blood was collected every Friday and cells were analyzed by Hemavet. Survival was monitored daily, and moribund mice were humanely sacrificed. Spleens were weighed at endpoint and normalized to mouse body weight at endpoint.

#### sAML patient-derived xenograft (PDX) models

PDX models were performed following protocol as previously described(32, 33). Briefly, PBMCs from sAML patients were isolated by Ficoll gradient extraction and CD34+ cells were isolated using magnetic enrichment (Miltenyi Biotec #130-100-453) and cultured overnight in SFEMII media (StemCell Technologies #09605) supplemented with Pen-Strep (50 Units/mL), human stem cell factor (SCF; 50 ng/mL), human thrombopoietin (TPO; 50 ng/mL), and human FLT3L (50 ng/mL). Human cells were transplanted into sublethally irradiated (200 cGy) 6-9 week-old NOD-scid-Il2rg-null-3/GM/SF (NSGS; The Jackson Laboratory #013062). NSGS mice were transplanted with 1 x 10^5^ cells via intra-tibial injection under anaesthesia with intraperitoneal injection of ketamine/xylazine mixture (2 mg/mouse; KetaVed). Four-to-five weeks after transplant, peripheral blood was collected and engraftment was evaluated by flow cytometry. Mice were treated with PBS vehicle intraperitonially, 5% dimethyl acetamide 0.5% methylcellulose by oral gavage, daily intraperitoneal BCI-HCl (25 mg/kg), ruxolitinib (90 mg/Kg) by oral gavage twice a day, or combination. Treatment began four weeks after transplant for sAML11 and after five weeks for sAML14, Monday through Friday. Mice were treated for four weeks and peripheral blood was collected every Friday and cells were analyzed by flow cytometry. Spleens and livers were weighed at endpoint and normalized to mouse body weight at endpoint.

### Publicly available databases accessed

Description and analyses of additional publicly available databases investigated in this study are as follows:

#### OHSU BeatAML

Clinical and expression data were obtained from Tyner *et al.*(*34*) and vizome.org. mRNA expression is indicated as normalized RPKM and samples compared were binned as NBM, prior-MPN sAML, prior-MDS sAML, and *de novo* AML.

#### GSE30029 *De novo* AML cohort(35)

Expression arrays of CD34+ and CD34- sorted mononuclear cells from AML patients and normal CD34+ bone marrow cells. mRNA expression is indicated as quantile normalized value.

#### GSE53482 primary myelofibrosis cohort(36)

Expression array of CD34+ from primary MF patients and CD34+ NBM. Gene expression value is indicated as log2 RMA signal. DEG analysis was performed by GEO2R comparing MF to NBM. For genes with multiple probes, the value with lowest adjusted *p*-value post-DEG was utilized.

#### TCGA LAML(37) and other TCGA datasets

Gene expression and mutational data was accessed from cBioportal(38) and pearson correlations were performed. Clinical data were extracted from source publication. Kaplan-Meier survival data of *RPS6KA1*-stratified patients and *log-rank p-value* were calculated, where event free survival and overall survival of the patients with the highest 50% expression (n = 86) were compared to the bottom 50% (n = 87).

For multivariate analysis, patient clinical information was retrieved from Supplemental Table 01 of TCGA LAML(37). *RPS6KA1* expression was extracted from cBioportal from the TCGA LAML cohort. A modified table encompassing clinical information and *RPS6KA1* expression was constructed for 173 patients and consisted of the following features: TCGA patient ID, *RPS6KA1* expression, Expired? 3.31.12, Sex, Race, FAB, Age, %BM blast, WBC, %PB blast, RISK (Cyto), RISK (Molecular), 2017 ELN Genetic Risk, OS months 3.31.12, Chemotherapy (pooled into ATRA, induction, hypomethylating, or other), Transplant (pooled into none, auto, or other), mutations/alterations occurring in at least 5 patients and pooled as present or null: *PML-RARA, MLL*-partner, *MYH11-CBFB, RUNX1-RUNX1T1*, *MLLT10*-partner, *BCR-ABL, GPR128-TFG, NUP98-NSD1, MLL-PTD, FLT3, NPM1, DNMT3A, IDH2, SRSF2, IDH1, RUNX1, TET2, TP53, NRAS, CEBPA, WT1, PTPN11, KIT, KRAS, MT-CO2, TTN, U2AF1, SMC1A, SMC3, STAG2, MT-CYB, PHF6, ASXL1, FAM5C, FCGBP, MUC16, RAD51*, # of mutated/altered genes, and mutated recurrent genes. Univariate Cox proportional hazard models were used to assess the effect of each factor on overall survival. Only those variables significant in the univariate analysis were considered for multivariate Cox model. The final Cox model was fitted using a backward selection procedure. The diagnosis of model assumptions was performed using residual plots and utilized 170 observations.

*RPS6KA1* expression data from the TCGA Pan-Cancer cohort was extracted from cBioportal. The number of patients (n =) from each cancer subtype is as follows: Acute Myeloid Leukemia (173), Adrenocortical Carcinoma (78), Bladder Urothelial Carcinoma (407), Brain Lower Grade Glioma (514), Breast Invasive Carcinoma (1082), Cervical Squamous Cell Carcinoma (36), Cholangiocarcinoma (36), Colorectal Adenocarcinoma (592), Diffuse Large B-Cell Lymphoma (48), Esophageal Adenocarcinoma (181), Glioblastoma (160), Head and Neck Squamous Cell Carcinoma (515), Kidney Chromophobe (65), Kidney Renal Clear Cell Carcinoma (793), Liver Hepatocellular Carcinoma (366), Lung Adenocarcinoma (510), Lung Squamous Cell Carcinoma (484), Mesothelioma (87), Ovarian Serous Cystadenocarcinoma (300), Pancreatic Adenocarcinoma (177), Pheochromocytoma and Paraganglioma (178), Prostate Adenocarcinoma (493), Sarcoma (253), Skin Cutaneous Melanoma (443), Stomach Adenocarcinoma (412), Testicular Germ Cell Tumors (149), Thymoma (119), Thyroid Carcinoma (498), Uterine Carcinosarcoma (57), Uterine Corpus Endometrial Carcinoma (527), and Uveal Melanoma (80). *RPS6KA1* expression is provided as RSEM (Batch normalized from Illumina HiSeq_RNASeq V2 (log2(value+1)).

#### Tzelepis *et al.* CRISPR dropout screen(39)

Dropout candidates in AML cell lines (such as *RPS6KA1*) at FDR 10% that were not essential in non-AML lines HT29 and HT1080 (#non-AML = 0) were identified and plotted.

#### *RPS6KA1* gene dependency from DepMap(40) and expression data from the Cancer Cell Line Encyclopedia (CCLE)(41)

Gene expression of DUSP family genes across AML cell lines, and *RPS6KA1* (Expression Public 21Q1) were accessed from CCLE by the Broad Institute. Gene Effect scores (CERES) from CRISPR (Avana) Public 21Q1 were accessed from the DepMap portal by the Broad Institute.

#### Cell line drug sensitivity correlations(42)

Fedratinib IC50 of AML cell lines were derived from Cancerxgene and the GDSC1 dataset, and correlated with CCLE *RPS6KA1* mRNA expression. Area under curve (AUC) of apitolisib, KW-7-42-1, torin-2, WYE-125132, MK-2206, and PF-4708671 and *DUSP6* expression were accessed from DepMap portal by the Broad Institute.

#### GSE153319 Serial scRNA-seq of CD34+ cells from primary MF patient(43)

Violin plots were constructed from individual barcoded cells across PMF, treatment PMF, and sAML timepoints, and relative fold change expression of *DUSP6*, *KLF2*, and *KLF1*, in addition to associated statistical analyses were derived from source publication.

#### GSE125150 inducible *KLF1*-inducible iPSC macrophages(44)

*p-values* of gene expression comparisons of *DUSP6* between untreated iKLF1.2 derived macrophages and tamoxifen-added iKLF1.2 derived macrophages were assessed.

#### GSE149119 murine bone marrow-derived macrophage *Klf2*-knockout(45)

*Dusp6* expression between vehicle and KO were compared with Mann-Whitney U-test

#### GSE27602 murine Klf2-/- yolk sac erythroid cells(46)

Multi-Chip Significance score (S-score) and expression comparison *p-*value of *Dusp6* (probe 1415834_at) between WT and Klf2-/- samples were identified. As per source publication, probe with absolute S-score values greater or equal to 2.00 were considered to be significant.

#### ChIP-seq

Sequencing tracks were investigated using the Cistrome Data Browser and WashU Browser with the following accession numbers: *Nfil3* (GSM1437733)(47); *Atf3* (GSM2663858)(48), *Fos* (GSM1875490(49), GSM2663847(48)), *Fosl2* (GSM1004808(50)), H3K27ac (GSM2974670(51), GSM851270(52)).

### Other statistical analysis

Statistical analyses were performed using GraphPad Prism (San Diego, CA) and *R* software. One-way ANOVA, two-way ANOVA, two-tailed Student’s t test, Mann Whitney U test, and Pearson correlations were performed as indicated. *, *p* < 0.05; **, *p* < 0.01; ***, *p* < 0.001; ****, *p* < 0.0001. All measurements were taken from distinct samples. All relevant assays were performed independently at least 3 times.

### Data availability

Patient microarray, bulk RNA-sequencing, scRNA-seq data, and mass cytometry data will be made publicly available upon publication. Analyzed sequencing data are currently presented in Table 2, 3, 4, 5, 7, and 8.

### Code availability

R scripts utilized in this study are available from corresponding author upon request.

## Results

### Transcriptome analysis reveals enriched proliferative signaling pathways and elevated *DUSP6* expression in secondary acute myeloid leukemia stem/progenitor cells

To understand alterations to the transcriptional landscape underlying MPN disease progression, we performed microarray analysis on sorted linage-negative (lin-) CD34+ HSPCs from 14 MF and 6 sAML patients, in addition to sorted CD34+ bone marrow cells from 5 healthy donors. We compared MF and sAML patient samples and plotted the most differentially expressed genes (DEGs) (Supplementary Fig. S1A, Table 1). Hallmark gene set enrichment analysis (GSEA) of the top altered pathways revealed enhancement of numerous pathways including TNF-⍺ signaling via NFκB, consistent with our previous findings(25, 27), in addition to interferon, IL6/JAK/STAT3, KRAS, IL2/STAT5, and PI3K/AKT/mTOR signaling in sAML vs MF HSPCs (Fig. 1A). Of the 457 statistically significant DEGs that were identified, notably, we observed enrichment of *IL3RA*, *FLT3*, and *CSF1R* (Fig. 1B) in sAML samples, which have been established as hyperactive disease drivers of *de novo* AML(53–55). These findings suggest shared cellular mechanisms underlying secondary and *de novo* AML. *DUSP6*, encoding a MAPK phosphatase that regulates ERK signaling, was also among the top DEGs and significantly elevated in sAML vs MF HSPCs (Fig. 1B, C), ranking higher than the only other DUSP family member identified, *DUSP18*, among the 23 DUSPs (Table 2).

**Figure 1.**
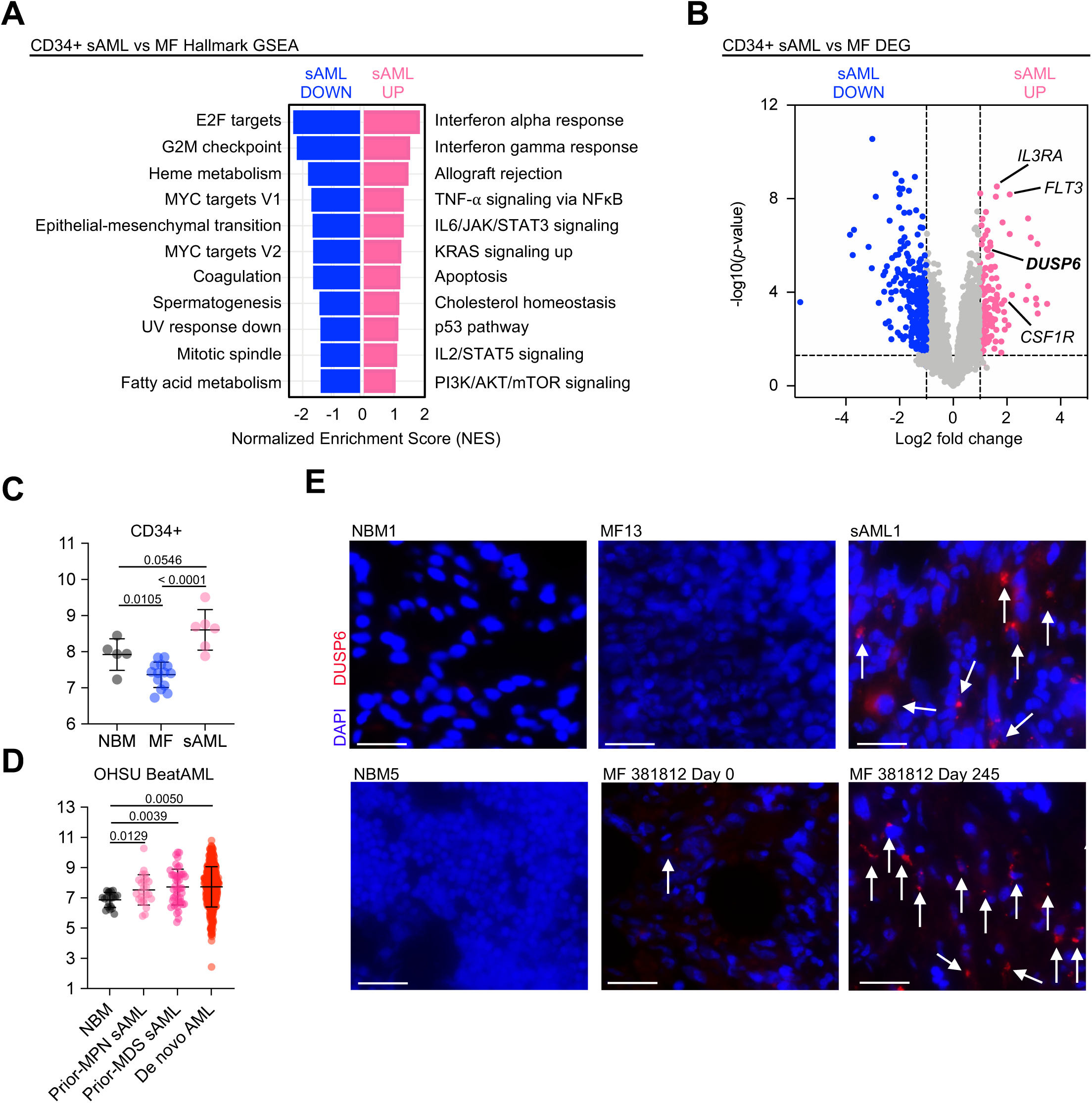
Elevated DUSP6 expression in patients with sAML compared to MPN. A) Gene set enrichment analysis of top Hallmark pathways enriched and diminished in linage-negative CD34+ cells from sAML patient samples (n = 6) compared to MF (n = 14) from microarray analysis. B) Volcano plot showing top elevated candidates in lineage-negative CD34+ cells from sAML patients compared to MF from microarray analysis. Dashed vertical lines denote boundaries of genes having at least 1 log2 fold change. Dashed horizontal line denotes p-value at 0.05 (−log10(p) = 1.3). C) *DUSP6* expression from CD34+ cells isolated from NBM (n = 5), MF (n = 14), and sAML (n = 6) patient samples. *DUSP6* values represent RMA from microarray. Statistics were assessed by two-tailed Student’s t test. D) *DUSP6* expression from healthy, individual bone marrow mononuclear cells (NBM MNC, n = 19), prior-MPN sAML MNC (n = 24), prior-MDS sAML MNC (n = 45), and *de novo* AML MNC (n = 375) from the OHSU BeatAML cohort. *DUSP6* values represent normalized RPKM. Statistics were assessed by two-tailed Student’s t test. E) Immunofluorescence of bone marrow from MF, sAML, and healthy donors. White arrows denote DUSP6 positive cell staining. Scale bar: 50 µM.

Aberrant upregulation of *DUSP6* expression in post-MPN sAML was corroborated by an independent database, the OHSU BeatAML cohort(34) (Fig. 1D), in which *DUSP6* was also enriched in other AML subtypes such as prior-MDS sAML and *de novo* AML, further suggesting a conserved role in AML development from different origins. Additionally, we found increased *DUSP6* in patient CD34+ AML cells compared to CD34+ NBM cells from a separate *de novo* AML cohort(35) (Supplementary Fig. S1B). Next, we confirmed elevated DUSP6 protein expression using immunofluorescence (IF) which revealed higher DUSP6 protein in bone marrow from sAML patients compared to MF and healthy donors (Fig. 1E, Supplementary Fig. S1C). Additionally, IF analysis of serial bone marrow from patient 381812 demonstrated a substantial increase in DUSP6+ cells at MF day 245 compared to day 0 (Fig. 1E). These findings were corroborated by imaging mass cytometry analysis revealing enrichment of CD34+ DUSP6+ cells from sAML1 compared to MF20 in addition to DUSP6 expression in subsets of monocytes, T cells, and B cells (Supplementary Fig. S1D). These data show that DUSP6 is upregulated in malignant blood cells, and during evolution of MPN to sAML.

### Single cell RNA sequencing of serial patient samples corroborates aberrant *DUSP6* expression accompanying disease progression

To understand patient-specific gene expression changes across disease evolution, we performed single cell RNA sequencing (scRNA-seq) in conjunction with TotalSeq surface protein marker detection on more than 50,000 sorted CD34+ cells of serial samples from three patients: patient 381812 at MF and sAML, patient 145790 at early MF, late MF, and sAML, and patient 374024 at PV and sAML– in addition to two NBM controls (N34, N39) (Fig. 2A). UMAP analysis showed tight clustering of cells from N34 and N39, whereas there was marked separation of MF and AML samples from 381812, and PV and sAML samples from 374024, suggesting distinct disease states (Fig. 2B). 145790 early MF clustered more closely to its late MF and AML stages, with the latter showing extensive overlap. Furthermore, there was proximal clustering between 381812 MF and 145790 early MF, and these MF stages were separated from 374024 PV. These observations support the notion that across patients there may be conserved similarities in HSPCs at an early MPN stage that branch into distinct entities during leukemic progression.

**Figure 2.**
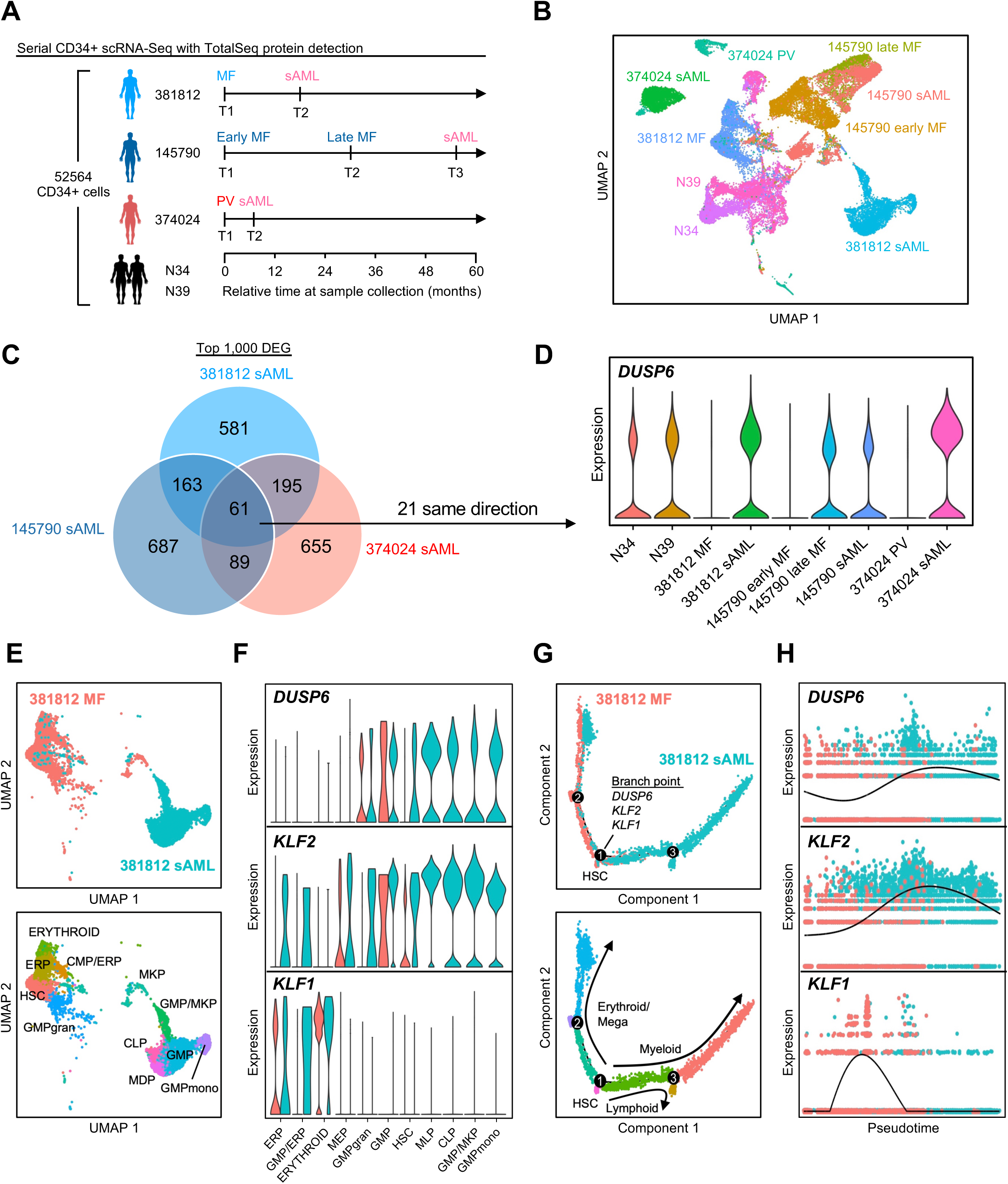
Serial patient CD34+ scRNA sequencing reveals elevated *DUSP6* along MPN to sAML progression. A) Schematic of scRNA-seq with Totalseq protein detection of 52,564 sorted CD34+ cells of serial samples from three MPN to sAML patients and two healthy bone marrow donors (N34, N39). Timeline shows disease stage and time (months) at which the sample was collected relative to early/first MPN stage. B) UMAP clustering of CD34+ cells from chronic MPN and sAML patients, and healthy donors. C) Venn diagram showing overlapping genes from the top 1,000 differentially expressed genes in serial CD34+ samples at the sAML stage compared to MPN stage of three patients (381812 MF sAML vs 38182 chronic MF, 145790 sAML VS 145790 early and late MF, 374024 sAML vs 374024 chronic PV). D) Violin plot showing elevated *DUSP6* expression at sAML stage compared to prior MPN stage. Bone marrow from two healthy donors (N34, N39) are included as references. E) UMAP analysis of MF and sAML disease states (top) and identified cell populations (bottom) in patient 381812. F) Violin plots of *DUSP6, KLF1,* and *KLF2* across identified subpopulations in 381812. G) Trajectory analysis showing differentially expressed effectors *DUSP6*, *KLF1*, and *KLF2* at identified branch points in MF and sAML disease states (top) and identified cell populations (bottom) from patient 381812. H) Pseudotime analysis of dynamic *DUSP6*, *KLF1*, and *KLF2* expression across disease progression from MF to sAML in patient 381812.

GSEA of the DEGs of the sAML stage to its paired chronic MPN showed TNF-⍺ signaling via NFκB to be the top pathway enriched across all three patients (Supplementary Fig. S2A). Of the top 1,000 DEGs from each paired sample, 61 were shared across all 3 patients and 21 were in the same direction (all sAMLs showed upregulation or downregulation) (Fig. 2C, Table 3). Within this shared group of 21 aberrantly expressed genes, *DUSP6* was elevated at the sAML stage from each of the three patients (Fig. 2D, Supplementary Fig. S2B). Notably, high *DUSP6* expression was also evident at the late MF stage from 145790. This observation of a rise in DUSP6 expression late in the chronic MPN stage, but prior to overt leukemic progression is consistent with the notion of dysregulated *DUSP6* expression as a driver, rather than simply a consequence, of disease progression.

Due to the intrinsic heterogeneity of sAML arising from antecedent PV vs MF, we further investigated the DEGs specifically between the two sAML evolved from MF (381812 and 145790), which shared 141 genes in the same direction (Supplementary Fig. S3A, Table 3). In addition to *DUSP6*, differential expression of KLF transcription factor family members was identified, notably the decreased expression of *KLF1* and increase of *KLF2*, which have been shown to play important roles in progenitor cell differentiation(56). Other transcription factors enriched included *NFIL3* (Nuclear factor, interleukin 3 regulated) in addition to its interacting partners including *ATF3*, *FOS*, and *FOSL2* that complex to form AP-1 and/or other multi-unit transcriptional regulators (Supplementary Fig. S3B). Publicly available ChIP-seq tracks(47–52) showed binding of these transcription factors to the *Dusp6* locus which also overlap with H3K27ac activation marks (Supplementary Fig. S3C), suggesting potential regulation of *Dusp6* by these canonical transcription factors. The family of early growth response (EGR) transcription factors was also enriched and has been described to demonstrate interplay with MAPK pathways(57). To substantiate these findings, we examined co-expression between *DUSP6* and these enriched genes of interest across our patient microarray and four additional publicly available patient cohorts(34–37). Overall, a consistent correlation of these genes with *DUSP6* was identified, with a negative correlation with *KLF1* and a significant positive correlation observed with *KLF2* that was significant across all five cohorts (Supplementary Fig. S3D). Supporting this, correlation analysis of single cells from 381812 showed *KLF2* to be the most positively correlated gene with *DUSP6*, while *KLF1* was among the 10 most negatively correlated genes (Supplementary Fig. S3E). These findings identify specific transcriptional programs as potential regulators of aberrant *DUSP6* expression across disease progression.

We then explored the cellular and transcriptional landscapes of MF and sAML stages in 381812 by first delignating major cell populations (Fig. 2E, Supplementary Fig. S4A, S4B). 381812 MF exhibited predominant enrichment of pluripotent HSCs, along with skewing of erythroid lineage populations, while 381812 sAML was largely comprised of more committed myeloid progenitors. We investigated if the elevated *DUSP6* expression at the sAML stage was indicative of high *DUSP6* in a distinct sAML subpopulation and/or if it was consistently elevated across all sAML clusters. We observed the latter, in sAML clusters having elevated *DUSP6* relative to MF clusters, and moreover, there was a similar pattern of enrichment of *KLF2* and suppression of *KLF1* among the dominant sAML clusters (Fig. 2F). Moreover, trajectory-based differential expression analysis revealed bifurcation of the MF and sAML samples, which overlapped with erythroid/megakaryocyte and myeloid identities, respectively, and identified *DUSP6*, *KLF2*, and *KLF1* among the significant branch point determinants (Fig. 2G, Table 4). Differentiation trajectory pseudotime analysis revealed concomitant elevation of *DUSP6* and *KLF2* and suppression of *KLF1* across state trajectories and disease progression from MF to sAML in 381812, which was further highlighted by similar trajectories in 374024 from PV to sAML and 145790 from early MF to late MF to sAML (Fig. 2H, Supplementary Fig. S5). In addition, we corroborated our findings in a separate serial CD34+ scRNA-seq dataset(43) of a PMF patient at different timepoints along transformation to sAML, in which we observed elevation of *DUSP6* and *KLF2,* in concomitant with decreased *KLF1* through chronic MPN to sAML progression (Supplementary Fig. S6A-C).

Further in support of our scRNA-seq findings, we also observed decreased *KLF1* expression and increased *KLF2* expression in sAML vs MF in our CD34+ patient microarray (Supplementary Fig. S6D). We sought to validate the functional effect of *KLF2* in the regulation of *DUSP6*, where its knockdown by shRNA in *JAK2* V617F mutant HEL cells led to downregulation of *DUSP6* mRNA and protein (Supplementary Fig. S6E S6F), and significantly inhibited proliferation (Supplementary Fig. S6G). Finally, publicly available databases revealed decreased *DUSP6* expression upon induction of *KLF1* in iPSC-derived macrophages(44), while *Klf2*-knockout in mouse bone marrow derived macrophages(45) and embryonic yolk sac erythroid cells(46) showed decreased *Dusp6* (Supplementary Fig. S6H, S6J). Altogether, bulk and single-cell sequencing analyses on primary patient samples demonstrate that *DUSP6* elevation is a marker of disease progression from MF to sAML, which may be regulated by a host of interplaying transcription factors including *KLF1* and *KLF2*.

### *DUSP6* inhibition suppresses JAK-STAT and S6 signaling pathways

Given the potential role of *DUSP6* in driving disease progression, we sought to further delineate its function in MPN/AML. From the Cancer Cell Line Encyclopedia (CCLE)(41), we first assessed the mRNA expression among DUSP family genes across 35 AML cell lines and observed high expression of *DUSP6*, which was also elevated in HEL cells (Supplementary Fig. S7A, S7B). Pharmacologic inhibition of DUSP6 via BCI led to a dose-dependent decrease in HEL cell viability (Supplementary Fig. S7C). At low concentrations, 24 hour BCI treatment increased ERK phosphorylation in HEL cells, consistent with the inhibitor’s capacity to suppress phosphatase activity (Supplementary Fig. S7D). Somewhat unexpectedly, higher BCI concentrations abolished ERK phosphorylation in conjunction with induction of apoptosis through increased cleavage of caspase 3. To determine if apoptotic and cell cycle inhibition profiles were a consequence of diminished ERK signaling, we treated HEL cells with the MEK1/2 inhibitor trametinib. Sub-micromolar doses of trametinib were sufficient to completely eradicate ERK phosphorylation but did not significantly alter cell viability or induce apoptosis (Supplementary Fig. S7C, S7D), suggesting that DUSP6 may play additional roles in signal transduction beyond direct ERK modulation to drive cancer survival.

To uncover novel effectors of DUSP6, we probed other predominant cancer signaling pathways and observed dose-dependent suppression of pS6 in addition to pSTAT3 and pSTAT5 in HEL cells by BCI treatment that was not observed with trametinib (Fig. 3A, Supplementary Fig. S7E, S7F). Further cell cycle and viability profiling showed BCI led to suppression of RB phosphorylation, induced G1 arrest, and induced apoptosis through increased annexin V staining (Fig. 3A-C). We also performed RNA-seq on HEL cells treated with 4 hours and 24 hours BCI, in which top downregulated GSEA pathways included E2F targets and G2M checkpoint (Fig. 3D, Supplementary Fig. S7G, Table 5), which were more pronounced with longer duration treatment. In addition to cell cycle related pathways, downregulation of MTORC1 signaling pathway was observed, consistent with signaling suppression in immunoblots. To confirm that the effects of BCI were DUSP6-dependent, genetic approaches were utilized to perturb *DUSP6* expression. Indeed, knockdown of *DUSP6* with independent shRNAs also led to suppression of pS6 and pSTAT3 in conjunction with reduced HEL viability (Fig. 3E, 3F). In contrast, ectopic expression of *DUSP6* increased cell viability, further suggesting an oncogenic role of *DUSP6* (Supplementary Fig. S7H, S7I). *DUSP6* overexpression also conferred greater resistance to BCI treatment compared to control vector (Supplementary Fig. S7J), further validating DUSP6 targeting via BCI.

**Figure 3.**
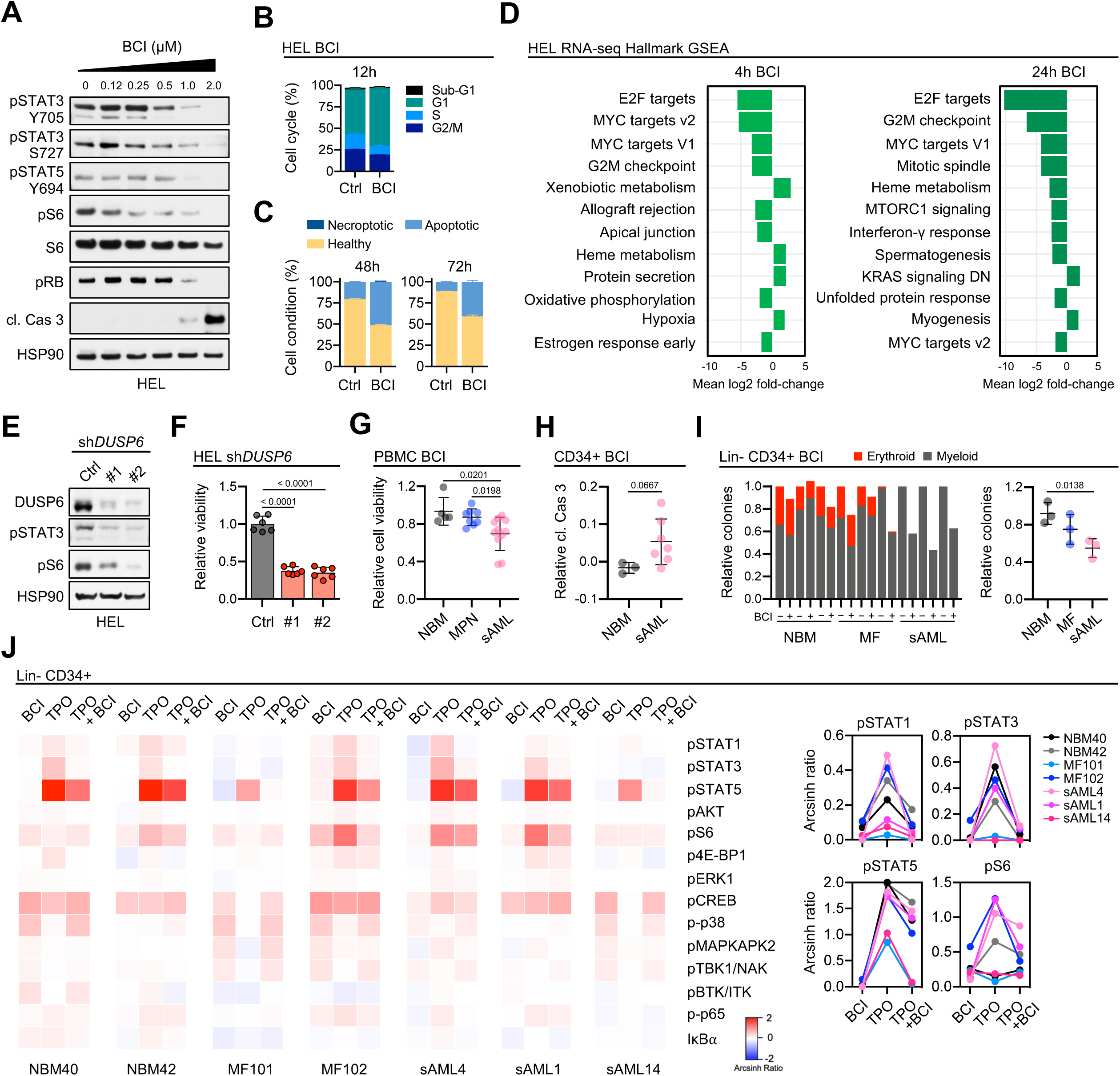
DUSP6 inhibition suppresses signaling from driver pathways and proliferation in MPN and AML. A) Immunoblot profiling of dominant signaling pathways in the *JAK2* V617F mutant HEL cell line. HEL cells were treated with increasing doses of DUSP6 inhibitor BCI for 24 hours. B) Cell cycle assay of HEL cells treated with 1µM of BCI or control for 12 hours. Cells were treated in triplicate (n = 3). C) Annexin V apoptosis assay of HEL cells treated with 1µM of BCI or control for 48 and 72 hours. Cells were treated in triplicate (n = 3). D) Hallmark gene set enrichment analysis of top altered pathways from RNA sequencing of HEL cells treated with 1µM BCI or DMSO control for 4 and 24 hours. E) Immunoblot of HEL cells upon knockdown of *DUSP6* with two independent shRNAs or pLKO control vector. F) Cell viability assay of HEL cells upon knockdown of *DUSP6* with two independent shRNAs or pLKO control vector. Cells were plated at n= 6 for each condition and grown for 96 hours with viability normalized to pLKO control vector. Mean and standard deviation presented. Statistics were assessed by two-tailed Student’s t test. G) Cell viability assay of normal donor bone marrow (NBM BMMC; n = 5), and PBMCs from MPN (n = 8), and sAML (n = 11) patients. All primary cells were from unique patients and were treated for 72 hours with 150 nM BCI. Statistics were assessed by two-tailed Student’s t test. H) Relative cleaved caspase 3 signal from CD34+ gated cells by mass cytometry. Unique sAML PBMCs (n = 7) and NBM BMMC (n = 3) were treated for 4 hours at 1 µM BCI. Arcsinh ratios indicate relative change in the BCI-treated condition compared to control. Statistics were assessed by two-tailed Mann-Whitney U test. I) Colony assay of sorted lin- CD34+ cells (n = 3 unique patients each for NBM, MF, and sAML). Sorted cells were grown in MethoCult H4034 Optimum for 12 days in 0.5 BCI µM or RPMI control. Samples were plated in duplicate. Statistics were assessed by Two-tailed Student’s t test. J) Heatmap and dot plots of altered signaling pathways of lin- CD34+ cells from unique NBM BMMCs, and PBMCs from MF, and sAML patients by mass cytometry. Patient samples were treated with 1 µM BCI for 4 hours, 20ng/mL TPO for 1 hour, or combination. Signals were normalized to the control treatment of each individual patient sample and reported as 90^th^ percentile Arcsinh ratio.

Consistent with the increased DUSP6 expression upon disease progression in patient samples, sAML peripheral blood mononuclear cells (PBMCs) were also more sensitive to BCI compared to MPN PBMCs and normal bone marrow mononuclear cells (BMMCs) (Fig. 3G), and BCI-treatment led to higher cleaved caspases 3 levels in CD34+ sAML cells compared to NBM (Fig. 3H). In lin- CD34+ colony formation assays, BCI significantly reduced the colony forming units from sAML samples, with smaller MF and sAML colonies observed compared to NBM, (Fig. 3I, Supplementary Fig. S7K). To better investigate BCI’s potency on distinct cell populations from primary samples and improve signal resolution to single cells, we performed multiplex profiling using suspension mass cytometry (CyTOF). Surface marker-guided dimensional clustering of sAML4 showed a large population of lin- CD34+ HSPCs, which demonstrated strong induction of pSTAT1, pSTAT3, pSTAT5, and pS6 upon treatment with JAK-STAT pathway activator thrombopoietin (TPO; Supplementary Fig. S8A, S8B), which was diminished by BCI co-treatment. Similar findings were observed in additional MF and sAML patient samples (Fig. 3J). Further mass cytometry analysis identified CD14+ monocytes as the predominant cytokine producing population, consistent with our previous studies(25) (Supplementary Fig. S8C). Upon TPO stimulation, IL-8 and in particular MIP-1β/CCL4 were strongly upregulated in CD14+ monocytes but were nearly completely abrogated with addition of BCI (Supplementary Fig. S8D). TPO-induced hyperproduction of MIP-1β/CCL4 in CD16+ monocytes and CD123+ cells from sAML5 were also suppressed by BCI (Supplementary Fig. S8E). Collectively, these novel findings demonstrate that *DUSP6* drives tumorigenicity in MPNs/sAML, and DUSP6 inhibition suppresses dominant pS6/STAT signaling pathways and suppresses inflammatory cytokine production.

### DUSP6 regulates S6 phosphorylation in MF and AML via indispensable RSK1

Alterations in PI3K/AKT/mTOR signaling pathways identified from bulk transcriptome and scRNA-seq analyses, in conjunction with observed suppression of pS6 after DUSP6 inhibition, led us to further explore downstream signaling mechanisms. Investigation of canonical regulators of S6 activity such as PI3K/AKT/mTOR and ribosomal S6 kinase (S6K) revealed a strong correlation between *DUSP6* and the area under curve (AUC) of inhibitors against these targets across multiple AML cell lines (Supplementary Fig. S9A). We hypothesized that these signaling nodes may be crucial to AML and subsequently explored a publicly available CRISPR dropout screen(39) which identified genes indispensable for survival of AML cell lines (MOLM-13, MV4-11, HL-60, OCI-AML2, and OCI-AML3) but not for non-AML control cell lines (HT29 and HT-1080). *RPS6KA1* (also known as ribosomal protein S6 kinase alpha-1, RSK1 or p90S6K) was among the 27 targets whose inhibition was lethal to at least 4 out of 5 AML lines, and was dispensable in both control lines (Supplementary Fig. S9B). RSK1 is the p90 subunit which together with beta-subunit p70S6K (*RPS6KB1*) forms S6K to phosphorylate S6. Consistent with these findings, we observed that *RPS6KA1* expression was highest in AML across a large collection of cancer cell lines from the Cancer Dependency Map Project (DepMap)(40) (Fig. 4A), and that AML was most sensitive to CRISPR-Cas9 mediated knockout of *RPS6KA1* compared to other cancers (where a lower CERES score is indicative of an essential gene, a score of 0 indicates non-essential, and a score of −1 is comparable to the median score of pan-essential genes; Figure 4B). Elevated expression of *RPS6KA1* was also observed in the TCGA Pan-Cancer cohort (10,071 total patients), in which patients with AML had the second highest expression out of 31 distinct cancer subtypes (Supplementary Fig. S9C).

**Figure 4.**
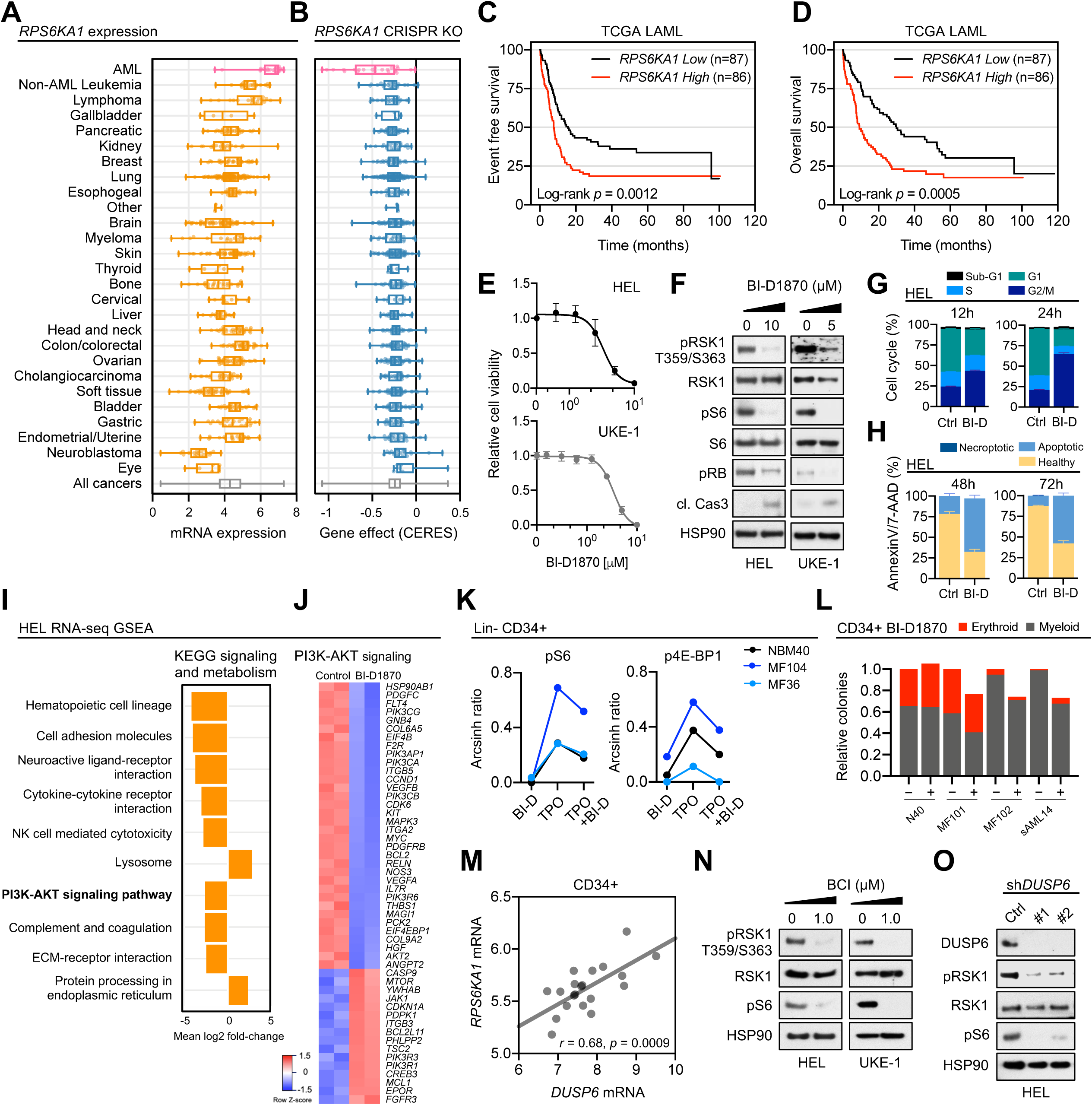
DUSP6 regulates S6 signaling through indispensable RSK1. A) Relative *RPS6KA1* expression across 804 pan-cancer cell lines from the Cancer Cell Line Encyclopedia. B) Gene dependency score for *RPS6KA1* CRISPR knock-out across 804 pan-cancer cell lines from the DepMap Portal. C) Kaplan-Meier event free survival curve of patients stratified by *RPS6KA1* expression from the TCGA LAML cohort (n = 173). D) Kaplan-Meier overall survival curve of patients stratified by *RPS6KA1* expression from the TCGA LAML cohort (n = 173) E) Cell viability curve of J*AK2* V617F mutant cell lines HEL and UKE-1 treated with increasing concentrations of RSK1 inhibitor BI-D1870. Cells were treated for 72 hours and viability was normalized to the control treatment. n = 6 replicates at each drug dose. Mean and standard deviation presented. F) Immunoblot of HEL and UKE-1 cells treated with BI-D1870 or control for 24 hours. G) Cell cycle assay of HEL cells treated with 5 µM BI-D1870 or control for 12 and 24 hours. Cells were treated in triplicate (n = 3). H) Annexin V apoptosis assay of HEL cells treated with 5 µM of BI-D1870 or control for 48 and 72 hours. Cells were treated in triplicate (n = 3). I) Gene set enrichment analysis showing top altered pathways from the KEGG signaling and metabolism gene set from RNA sequencing of HEL cells treated with 10 µM of BI-D1870 or control for 24 hours. J) Top 50 differentially expressed genes from the KEGG PI3K-AKT signaling pathway of RNA-seq from HEL cells treated with 10 µM of BI-D1870 for 24 hours vs control treatment. K) Dot plots of altered signaling pathways of lin- CD34+ cells from unique normal bone marrow donors and peripheral blood of MF patients by mass cytometry. Patient samples were treated with 5 µM BI-D1870 for 4 hours, 20ng/mL TPO for 1 hour, or combination. Signals were normalized to the control treatment of each individual patient sample and reported as 90^th^ percentile Arcsinh ratio. L) Colony assays of sorted lin- CD34+ cells from unique normal bone marrow, MF and sAML patients. Sorted cells were grown in MethoCult H4034 Optimum for 12 days in 1.5 µM BI-D1870 or RPMI control. Samples were plated in duplicate. M) Correlation of *DUSP6* and *RPS6KA1* from lin- CD34+ cells from MF (n=14), and sAML (n=6) patient samples. Expression values represent RMA from microarray. N) Immunoblot of HEL and UKE-1 cells treated with 1 µM BCI or control for 24 hours. O) Immunoblot of HEL cells after knockdown with two independent shRNAs targeting *DUSP6* or non-targeting pLKO control

In keeping with the dependency of RSK1 in AML cells, Kaplan-Meier event free survival and overall survival analysis of AML patients from TCGA showed significantly worse outcome for AML patients with higher *RPS6KA1* expression (Fig. 4C, 4D). Multivariate analysis using Cox proportional-hazards model on overall survival showed *RPS6KA1* expression to be significant (HR= 1.60, 95% CI: 1.10, 2.34) and comparable to other important factors: age (HR = 1.04, 95% CI: 1.02, 1.05), WBC (HR = 1.01, 95% CI: 1.00, 1.01), molecular risk classification of poor (HR = 2.52, 95% CI: 1.29, 4.94), alterations in *TP53* (HR = 2.99, 95% CI: 1.49. 6.00), *RUNX1* (HR = 1.99, 95% CI: 1.10, 3.60), and *FLT3* (HR = 1.92. 95% CI: 1.23, 2.99; Table 6). These findings demonstrate that aberrant *RPS6KA1* expression is associated with poor prognosis in AML and indicate that RSK1 represents a candidate target for therapeutic intervention.

Treatment of HEL and a second *JAK2*-mutant leukemia line UKE-1 with RSK1/2/3/4 inhibitor BI-D1870 led to a dose-dependent decrease in cell viability concurrent with reduced phosphorylation of pRSK1 at key activation residues T359/S363 and the suppression of pS6 (Fig. 4E, 4F). Importantly, *RPS6KA1* knockdown phenocopied signal suppression seen with pan-RSK inhibition and significantly reduced HEL viability (Supplementary Fig. S9D, S9E), consistent with the AML CRISPR-KO phenotype (Fig. 4B). BI-D1870 treatment also inhibited cell cycle progression by arresting cells in G2 phase and led to increased cellular apoptosis (Fig. 4F-H). In addition, RNA-seq analysis of HEL cells treated with BI-D1870 for 24 hours or DMSO control revealed downregulation of PI3K-AKT signaling among the top altered GSEA KEGG signaling and metabolism pathways upon RSK1 inhibition (Fig. 4I, Table 7). Suppression of mediators of G2/M transition including *WEE1, PLK1*, *CDK1*, and *CCNA2* (Table 7) was also observed in BI-D1870 treated cells, and the top 50 DEGs in the PI3K-AKT signaling pathway showed suppression of other cell cycle G1/S phase mediators *CCND1* and *CDK6* and upregulation of *CDKN1A* (Fig. 4J). In accordance with our annexin V assay, downregulation of apoptosis regulators *BCL2*, and upregulation of *BCL2L11* and smaller isoform *MCL1* were also observed upon BI-D1870 treatment. Mass cytometry of MF patient samples treated with BI-D1870 showed inhibition of TPO-induced pS6 and p4E-BP1 signaling in lin- CD34+ HSPCs, but minimal effect on phosphorylation of STAT proteins, suggesting BI-D1870 largely targets downstream of mTOR (Fig. 4K, Supplementary Fig. S9F). BI-D1870 also preferentially suppressed formation of myeloid colonies from MF and sAML CD34+ HSPCs compared to normal bone marrow (Fig. 4L). These findings establish *RPS6KA1* as a novel biomarker predicting patient prognosis and an essential candidate in MPN/AML to be exploited therapeutically.

We observed similar elevated expression of *RPS6KA1* at single cell resolution in CD34+ cells from 381812 sAML compared to MF (Supplementary Fig. S9G), and in bulk CD34+ sAML patient cells, where a strong correlation with *DUSP6* was also noted (Fig. 4M, Supplementary Fig. S9H). Given this observed relationship, we sought to determine if suppression of pS6 after DUSP6 perturbation was mediated through RSK1 inhibition. BCI treatment in HEL and UKE-1 cells led to reduced T359/S363 phosphorylation of RSK1, which was consistently suppressed in *DUSP6* knockdown cells (Fig. 4N, 4O). In contrast, *RPS6KA1* overexpression conferred resistance to BCI (Supplementary Fig. S9I). Therefore, we explored combination therapy with the addition of BCI to sub-maximal doses of BI-D1870, which suppressed HEL and UKE-1 viability more than each agent alone (Supplementary Fig. S9J). GSEA of RNA-seq of HEL cells treated with both BCI and BI-D1870 showed amalgamation of altered pathways observed in monotherapy, demonstrating suppression of JAK/STAT and MTORC1 signaling pathways, E2F targets, and increased p53 pathway compared to DMSO control (Supplementary Fig. S9K). Mass cytometry analysis of lin- CD34+ cells from primary sample MF103 revealed suppression of pS6, pSTAT1/3/5, and p4E-BP1 proteins with BCI and BI-D1870 treatment, demonstrating combined signal-suppressive phenotypes from each inhibitor (Supplementary Fig. S9l). Taken together, these studies characterize a novel DUSP6-RSK1-S6 axis that is prognostic of patient outcome and of which perturbations in this cascade potently suppress MPN/sAML.

### DUSP6 inhibition overcomes persistence to fedratinib *in vitro* and *ex vivo*

Prolonged suppression of driver pathways in cancer through single agent therapy inevitably leads to resistance and selection of drug-persistent clones, thus requiring further understanding of these dynamic alterations to amend therapeutic strategies. Cancer persistence is observed in the clinic, as JAK2-targeted therapies fail to eradicate the underlying MPN malignant clone, rendering MPN patients susceptible to leukemic transformation. We sought to mimic JAK2 inhibitor persistence *in vitro* via prolonged culturing of HEL cells in the presence of either ruxolitinib or fedratinib to generate ruxolitinib-persistent (Rux-P) and fedratinib-persistent (Fed-P) cells lines (Fig. 5A). RNA-seq analysis revealed upregulation of *DUSP6* in Rux-P cells and Fed-P cells compared to parental HEL cells (Fig. 5B, Table 8). We also observed a relative increase of expression in a curated gene set of “*DUSP6* targeted genes upregulated”(58) in Fed-P cells (Fig. 5C). Ectopic *DUSP6* expression mediated resistance to both fedratinib and ruxolitinib in parental HEL cells (Fig. 5D), and upon JAK2 inhibition these cells retained pS6 but not pSTAT3, implicating pS6 as a key effector mediating resistance to JAK2 inhibitors (Fig. 5E). Notably, *DUSP6* overexpression rescued pSTAT3 despite BCI treatment, suggesting indirect modulation of STAT signaling via BCI. In contrast, knockdown of *DUSP6* further sensitized parental HEL cells to fedratinib and ruxolitinib (Fig. 5F, 5G). We also observed a trend of elevated *DUSP6* expression correlating with decreased sensitivity to fedratinib in 18 AML cell lines (Supplementary Fig. S10A). These findings highlight a pivotal role for *DUSP6* in mediating resistance to JAK2 inhibitors in AML.

**Figure 5.**
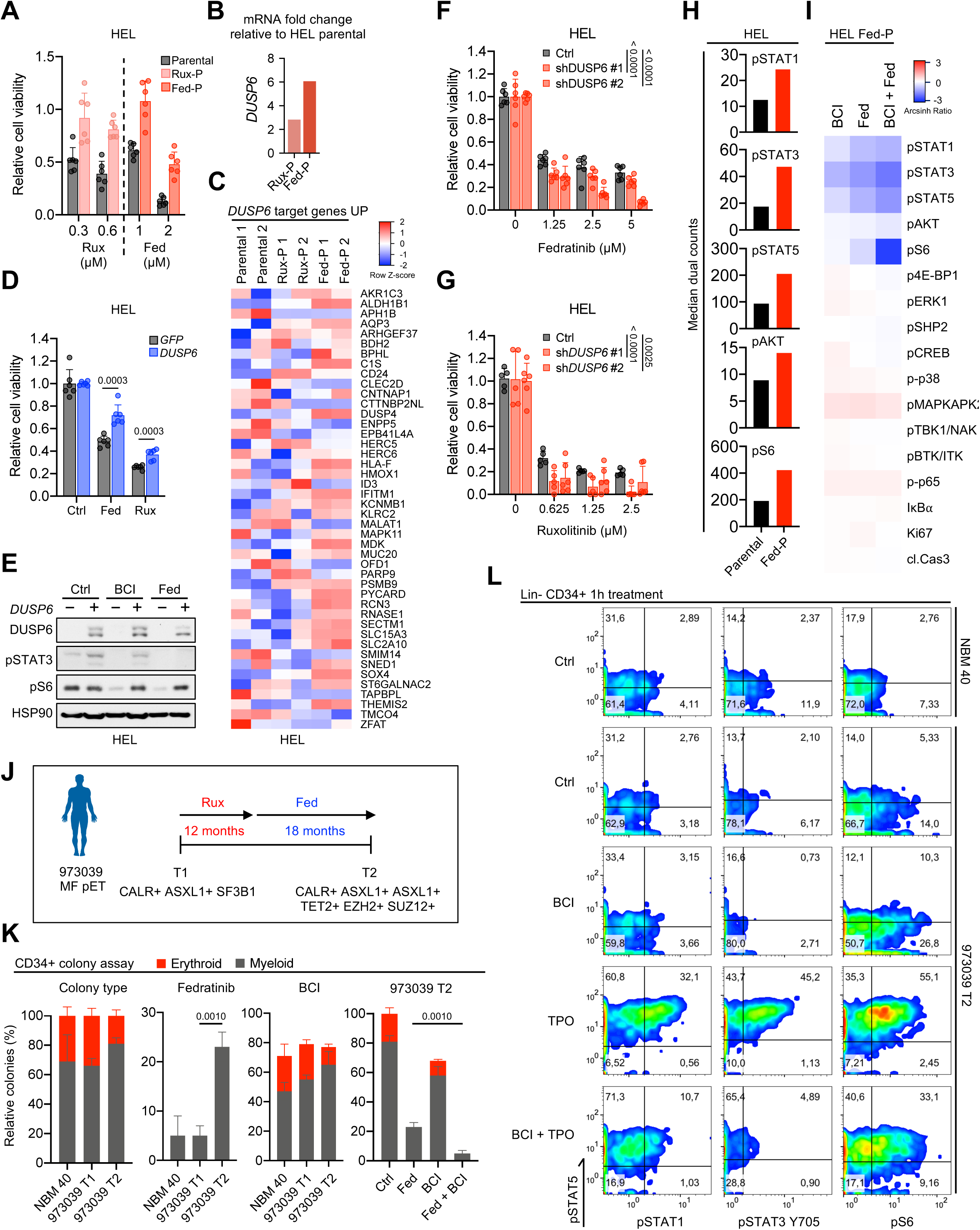
DUSP6 inhibition overcomes persistence to JAK inhibitors. A) Cell viability assay of parental HEL, HEL ruxolitinib-persistent (Rux-P), and HEL fedratinib-persistent (Fed-P) cells treated at indicated doses of JAK2 inhibitors ruxolitinib and fedratinib. Cells were treated for 72 hours and viability was normalized to the control. n = 6 replicates at each drug dose. Mean and standard deviation presented. B) Fold change of *DUSP6* mRNA expression in HEL persistent cells relative to HEL parental cells from RNA-seq analysis. C) Heatmap of a curated gene set of “*DUSP6* target genes UP” from RNA-seq analysis of HEL parental, HEL Rux-P, and HEL Fed-P cells. D) Cell viability assay of HEL parental cells treated with fedratinib and ruxolitinib after ectopic expression of *DUSP6* or a control vector. Cells were treated with 0.625 µM of JAK2 inhibitors for 96 hours and viability was normalized to the control treatment group from each population. n = 6 replicates. Mean and standard deviation presented. Statistics were assessed by two-tailed Student’s t test. E) Immunoblot of HEL cells ectopically expressing *DUSP6* or *GFP* control vector. Cells were treated with control, 1 µM BCI, or 1 µM fedratinib for 24 hours. F) Cell viability assay of HEL parental cells treated with fedratinib after knockdown with two independent shRNAs targeting *DUSP6* or non-targeting pLKO control. Cells were treated for 96 hours and viability was normalized to the control treatment group from each population. n = 5-6 replicates at each drug dose. Mean and standard deviation presented. Statistics were assessed by two-way ANOVA with Dunnett’s multiple comparisons test. G) Cell viability assay of HEL parental cells treated with ruxolitinib after knockdown with two independent shRNAs targeting *DUSP6* or non-targeting pLKO control. Cells were treated for 96 hours and viability was normalized to the control treatment group from each population. n = 6 replicates at each drug dose. Mean and standard deviation presented. Statistics were assessed by two-way ANOVA with Dunnett’s multiple comparisons test. H) Expression profiling of key phospho-proteins in HEL parental and HEL Fed-P cells by mass cytometry. Expression denoted as median dual counts. I) Mass cytometry heatmap of HEL Fed-P cells treated with BCI, fedratinib, or combination. Cells were treated with 1 µM BCI and/or 1 µM fedratinib for 1 hour and signals were normalized to the control treatment and reported as median Arcsinh ratio. J) Schematic of disease progression and time course of clinical treatments of patient 973039. K) Colony assays of sorted lin- CD34+ cells from 973039 at two timepoints, and healthy donor bone marrow (NBM 40). Sorted cells were grown in MethoCult H4034 Optimum for 2 weeks in the presence of inhibitors: Fedratinib panel: 1 µM fedratinib; BCI panel: 0.5 µM BCI; 973039 T2 panel: 1 µM fedratinib, 1 µM BCI, and 1 µM fedratinib + 1 µM BCI. Samples were plated in triplicate. Statistics were assessed by two-tailed Student’s t test. L) Paired mass cytometry analysis of lin- CD34+ cells from 973039 T2 and NBM 40. Cells were treated with 1 µM BCI for 1 hour, 20ng/mL TPO for 15 minutes, or combination.

Notably, Fed-P HEL cells showed similar sensitivity to BCI compared to the parental line (Supplementary Fig. S10B), while the addition of fedratinib to BCI also further suppressed cell viability in HEL parental cells (Supplementary Fig. S10C). Mass cytometry analysis of intracellular signaling changes in Fed-P cells revealed elevated phosphorylation of STAT1/3/5, AKT, and S6, relative to the parental line (Fig. 5H). BCI or fedratinib alone suppressed signaling through these pathways with combination treatment demonstrating enhanced inhibition, particularly of pS6 (Fig. 5I). Taken together, these findings demonstrate that BCI alone or in combination with fedratinib can effectively inhibit cell proliferation through suppression of JAK-STAT and S6 signaling in both naïve and fedratinib-persistent cells.

To extend these findings *ex vivo*, we examined serial samples from MF post-ET patient 973039 obtained prior to initiation of JAK2 inhibitor treatment (T1) (Fig. 5J). Upon poor clinical response to ruxolitinib, patient 973039 was switched to fedratinib treatment for 18 months with subsequent development of worsening anemia, a hallmark of disease progression and poor response to therapy, at which timepoint an additional sample was collected (T2). Consistent with disease progression, targeted sequencing revealed acquisition of *TET2*, *EZH2*, and *SUZ12* mutations, in addition to a second *ASXL1* mutation, at T2. We then isolated the CD34+ population and performed colony assays. 973039 T2 showed relatively decreased erythroid colonies and increased myeloid colonies compared to 973039 T1 and normal bone marrow (Fig. 5K), consistent with the clinical observation of worsening anemia. CD34+ cells from the later T2 timepoint were less susceptible than normal or T1 cells to inhibition of colony formation in the presence of fedratinib. Fedratinib also preferentially reduced erythroid colonies in all three samples, whereas BCI treatment reduced overall colony formation to a similar degree. Notably, the addition of BCI to fedratinib suppressed the residual, persistent colonies from the fedratinib-alone treatment in 973039 T2. Paired mass cytometry analysis of 973039 T2 showed that BCI was sufficient to reverse TPO induction of pSTAT1, pSTAT3, pSTAT5, and pS6 (Fig. 5I). In sum, these data indicate that DUSP6 targeting via BCI, either alone or in combination with fedratinib, represents a potentially effective and novel therapeutic avenue to overcome both naïve and JAK2 inhibitor-persistent MPNs.

### *DUSP6* inhibition suppresses disease burden and progression across multiple MPN and sAML mouse models

We then assessed the functional impact of DUSP6 targeting *in vivo* across MPN and sAML murine models. We first utilized *Jak2* V617F knock-in mice, which exhibit features of PV including erythrocytosis, leukocytosis, and splenomegaly(59). Mice transplanted with *Jak2* V617F-mutant cells were treated with vehicle or 25mg/kg BCI 5 days on, 2 days off (Fig. 6A). Hematocrit remained abnormally elevated in vehicle-treated control mice but was significantly reduced within the first week of BCI administration and sustained across the entire treatment duration (Fig. 6B). White blood cell (WBC) counts, individual WBC differentials, and platelet counts were similarly significantly suppressed in BCI-treated mice compared to vehicle-treated controls (Fig. 6B, Supplementary Fig. S11A). There was no significant difference in body weight in treatment groups, whereas BCI reduced splenomegaly and corrected liver weights (Fig. 6B, Supplementary Fig. S11B). Importantly, BCI treatment of primary wildtype mice did not pathologically cause anemia or other cytopenias, or decrease spleen and liver weights (Supplementary Fig. S11C), suggesting potency and specificity of BCI against MPN disease compared to normal tissue and confirming BCI does not exert non-specific myelosuppressive effects.

**Figure 6.**
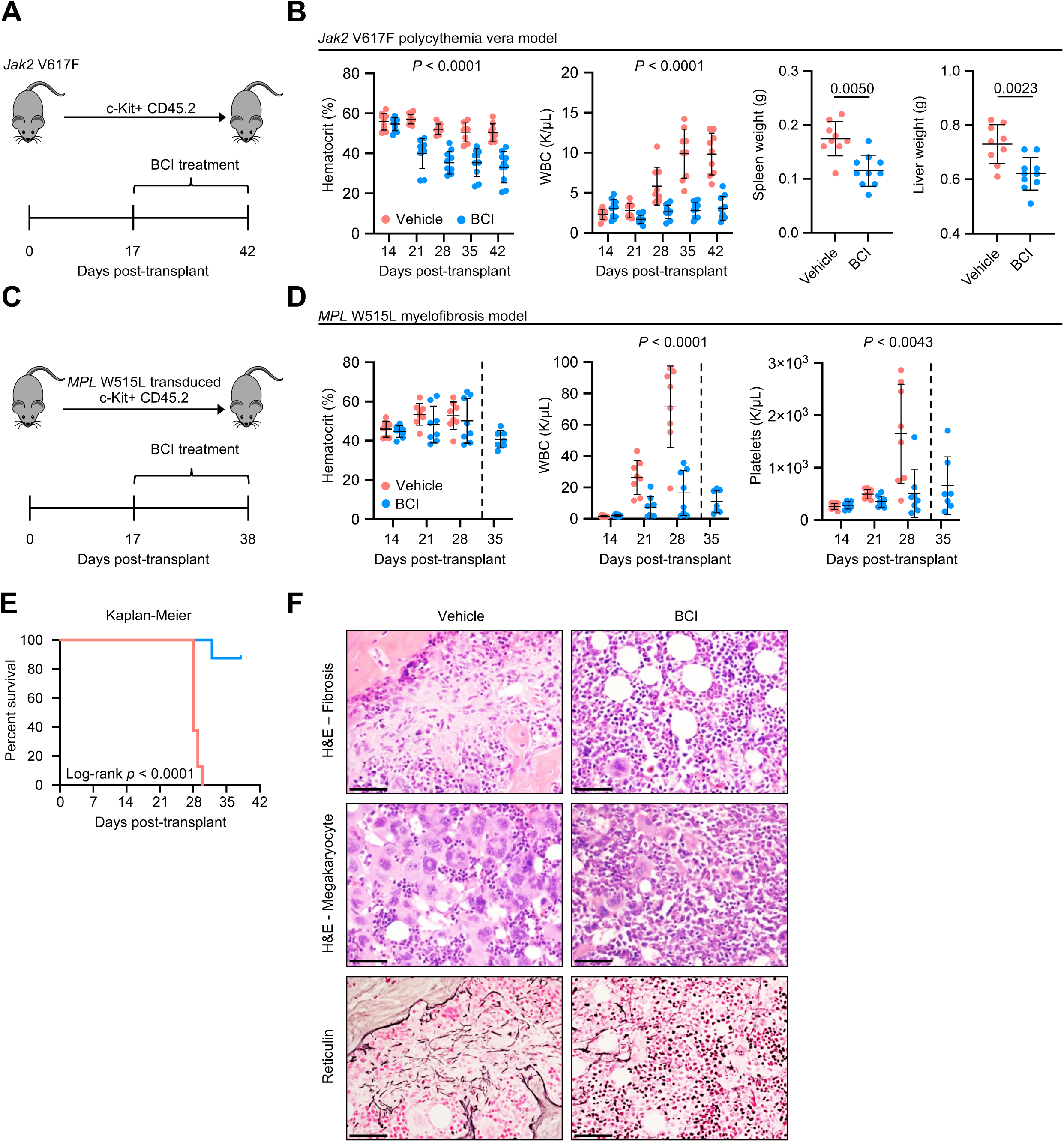
BCI is efficacious in MPN mouse models. A) Schematic of the *Jak2* mouse model. c-KIT+ cells from CD45.2 C57BL/6J mice were isolated and transplanted into CD45.1 C57BL/6J mice. Mice were treated with vehicle (n = 9) or 25 mg/kg BCI (n = 10) daily for 5 days followed by 2 days off treatment starting on day 17. B) Efficacy of BCI in the *Jak2* mouse model. Plots show hematocrit and white blood cell (WBC) counts of transplanted mice treated with vehicle or BCI across multiple timepoints and and spleen and liver weights at endpoint. Hematocrit and WBC counts statistics were assessed by two-way ANOVA comparing vehicle and BCI. Spleen and liver weights statistics were assessed by two-tailed Student’s t test. C) Schematic of the *MPL* mouse model. Retroviral *MPL* was transduced into c-KIT+ cells from CD45.2 C57BL/6J mice and transplanted into CD45.1 C57BL/6J mice. Mice were treated with vehicle (n = 8) or 25 mg/kg BCI (n = 8) daily for 5 days followed by 2 days off treatment starting on day 17. D) Efficacy of BCI in the *MPL* mouse model. Plots show hematocrit, WBC, and platelet counts of transplanted *MPL* mice treated with vehicle or BCI across multiple timepoints. WBC and platelet counts statistics were assessed by two-way ANOVA comparing vehicle and BCI. E) Kaplan-Meier survival analysis of transplanted mice treated with vehicle or BCI were assessed by log-rank test. F) Images show representative H&E and reticulin staining of bone marrow of tibias from vehicle and BCI treated *MPL* mice at endpoint. Scale = 50 µM.

We next evaluated the efficacy of BCI in the *MPL* W515L model, which confers a robust MF phenotype with marked leukocytosis, bone marrow fibrosis, and early lethality(60). Kit+ cells from CD45.2 mice were transduced with *MPL* W515L retrovirus and transplanted into lethally-irradiated CD45.1 recipient mice (Fig. 6C, Supplementary Fig. S11D). Mice receiving 25 mg/kg BCI showed no difference in hematocrit compared to vehicle but had significant reduction of leukocytosis and near significant reduction of normalized spleen weight (Fig. 6D, Supplementary Fig. S11E). Pathological thrombocytosis was also significantly reduced by BCI (Fig. 6D). While vehicle-treated mice were all moribund by day 30, BCI-treated mice exhibited markedly prolonged survival (Fig. 6E). Bone marrow H&E staining at endpoint demonstrated evident suppression of fibrosis and megakaryocyte hyperplasia in BCI-treated mice (Fig. 6F). Complementary immunohistochemical staining (IHC) showed reduction of reticulin with BCI treatment (Fig. 6F). Taken together, these findings demonstrate that BCI is efficacious in reducing disease burden of both PV and MF MPN mouse models.

To further recapitulate human disease, we adopted a novel, patient-derived xenograft (PDX) humanized animal system in which engrafted cells propagate MPN features(32, 33) (Fig. 7A). CD34+ cells from sAML post-MF 718407 (sAML11) were transplanted into NSGS mice, and engrafted mice were treated with 25mg/kg BCI or vehicle. BCI treated mice showed marked reduction in percentage of human CD45 (hCD45) cells in both peripheral blood and bone marrow (Fig. 7B). There was reversal of splenomegaly but no changes to liver weights at end point (Fig. 7B)

**Figure 7.**
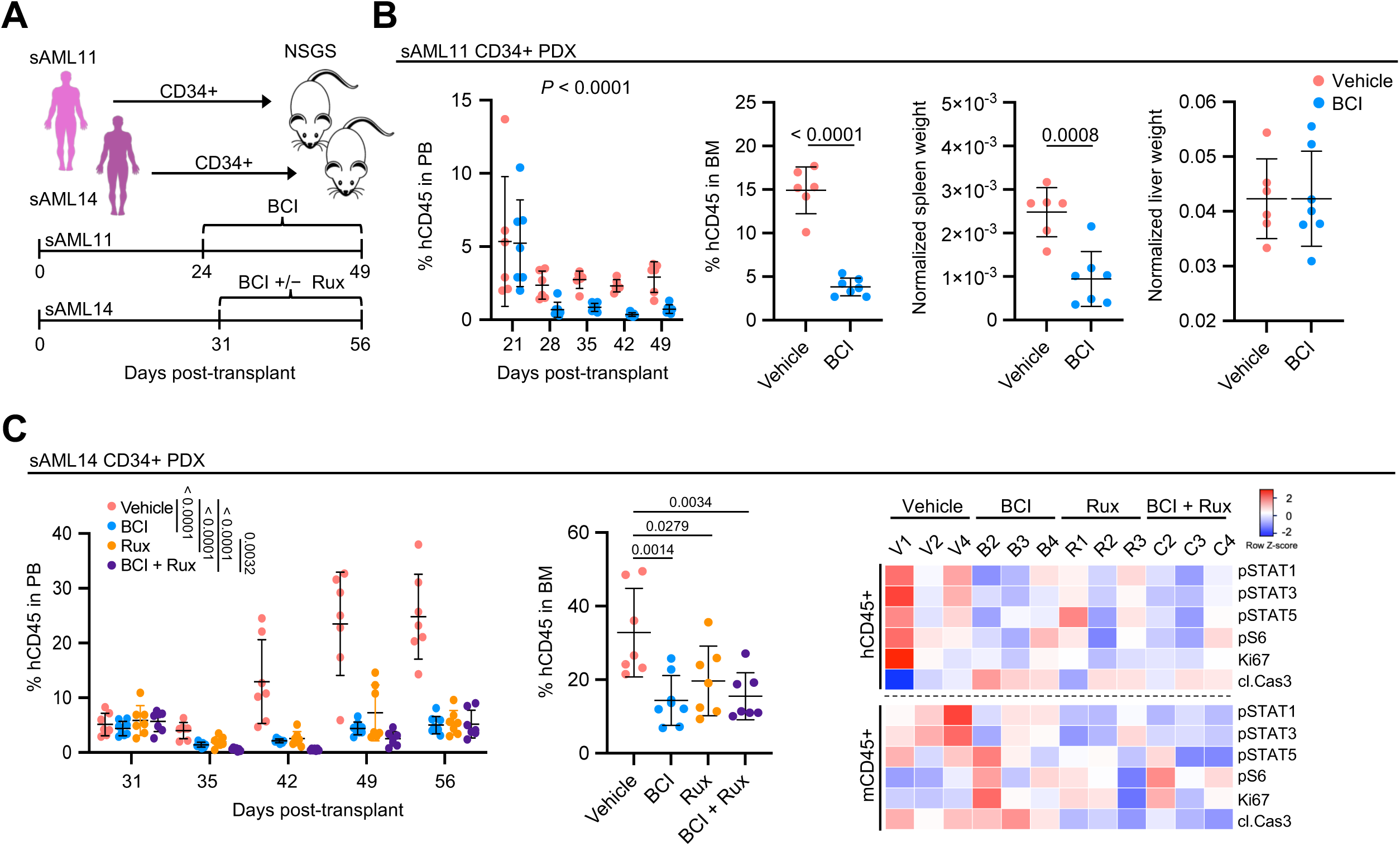
BCI alleviates disease burden across sAML patient-derived xenograft models. A) Schematic of the CD34+ sAML patient-derived xenograft mouse models. CD34+ cells were isolated from patients sAML11 and sAML14 and transplanted into NSGS mice. For the sAML11 PDX, mice were treated with vehicle (n = 6) or 25 mg/kg BCI (n = 7) daily for 5 days followed by 2 days off treatment starting on day 24. For the sAML14 PDX, mice were treated with vehicle (n = 7), 25 mg/kg BCI (n = 8), 90mg/kg ruxolitinib (n = 7), or combination (n = 7) following weekly schedule of 5 days on, 2 days off treatment starting on day 31. B) Efficacy of BCI in the sAML11 PDX model. Plots show percentage of human CD45 (hCD45) in peripheral blood and bone marrow of transplanted mice treated with vehicle or BCI across multiple timepoints and spleen weights of mice at endpoint normalized by mouse weight. %hCD45 in PB statistics assessed by two-way ANOVA comparing vehicle to BCI. %hCD45 in BM, and normalized spleen and liver weights statistics were assessed by two-tailed Student’s t test. C) Efficacy of treatment in the sAML14 PDX model. Plots show percentage of hCD45 cells in peripheral blood and bone marrow of transplanted mice treated with vehicle, BCI, ruxolitinib, or combination across multiple timepoints. %hCD45 in PB statistics were assessed by two-way ANOVA comparing vehicle vs each individual treatment group with Dunnett’s multiple comparison test, and two-way ANOVA for ruxolitinib vs combination. %hCD45 in BM statistics were assessed by one-way ANOVA with Dunnett’s multiple comparison test. Rightmost panel: Mass cytometry of PBMCs isolated from the bone marrow of three sAML14 PDX mice from each treatment group at endpoint. Heatmap denotes row z-score calculated from mean signal intensity of each marker.

We utilized a second PDX model engrafted with CD34+ cells from post-ET/MF sAML post-MF 784981 (sAML14) to explore efficacy of combination therapy with ruxolitinib and BCI. Similar to PDX sAML11, potent suppression of PB hCD45 engraftment was observed with BCI monotherapy compared to vehicle (Fig. 7C). Ruxolitinib treatment also reduced PB hCD45 engraftment, with combination treatment (BCI + ruxolitinib) further suppressing PB hCD45 engraftment compared to ruxolitinib alone. BCI, ruxolitinib, and combination treatment also all significantly suppressed BM hCD45 engraftment compared to vehicle and showed a trend in reduction of spleen but not liver weight (Fig. 7C, Supplementary Fig. S11F). To assess the degree of signaling suppression across the four treatment conditions, we performed mass cytometry on flushed bone marrow cells from sAML14 PDX mice (Supplementary Fig. S11G). BM hCD45+ cells isolated from BCI, ruxolitinib, and BCI + ruxolitinib treatment groups showed relative inhibition of pSTAT1/3/5, pS6, and Ki67 compared to vehicle, in conjunction with increased cleaved caspase 3 (Fig. 7C). Mouse BM hCD45+ cells (mCD45+), also showed suppression of STAT phosphorylation but no obvious trend in pS6, Ki67, or cleaved caspase 3. These experiments functionally validate signal suppression *in vivo* and demonstrate selectivity of BCI and ruxolitinib alone or in combination against engrafted malignant cells compared with normal tissue. Thus, our findings demonstrate a consistent reduction in disease burden across humanized and MPN mouse models, and highlight DUSP6 inhibition as a promising therapeutic approach.

## Discussion

Using bulk expression profiling and single-cell sequencing approaches on serial MPN and sAML patient samples, we identify aberrant overexpression of *DUSP6* and downstream effector *RPS6KA1* driving treatment resistance and disease progression. We target these dependencies using small molecule inhibitors BCI and BI-D1870, and through mass cytometry multiplex profiling highlight suppression of dominant STAT and S6 proliferation pathways across cell lines and primary samples. Finally, we demonstrate selective efficacy of BCI across *in vivo* models of PV and MF, and show that BCI treatment reduces disease burden alone and potentiates ruxolitinib efficacy in sAML PDX models.

Clinical observations of JAK2 inhibitors alone failing to eradicate the underlying malignant clone and limited anti-proliferative effects of MEK inhibitor trametinib alone suggest single pathway suppression to be insufficient in reducing disease in MPNs. Compensatory mechanisms of reduced therapeutic efficacy likely include both adaptive upregulations of driver pathways such as increased levels of pSTAT1/3/5, pAKT, and pS6 effectors seen in HEL Fed-P cells as well as elevated intrinsic signaling driving disease transformation from chronic MPN phase to sAML. Recently, the merits of combinational therapy have been explored, with emphasis on concurrent inhibition of JAK/STAT and MAPK pathways. The addition of trametinib to ruxolitinib suppressed MEK/ERK feedback activation in *Jak2* V617F and *MPL* W515L mice and reversed bone marrow fibrosis greater than in monotherapy(12). Similar therapeutic strategies also reduced disease penetrance and *JAK2* V617F clonal burden in MPN xenograft models(11). Our findings with BCI indicate that DUSP6 targeting impacts MAPK in addition to multiple downstream signaling effectors. Continued investigation utilizing additional screening or synthetic lethal approaches may distinguish essential targets for rational therapeutic intervention.

Current understanding of disease transformation from chronic MPN to sAML is complicated by relationships between distinct cellular populations, diverse genomic alterations and temporal interactions. Our sequencing approaches led us to identify a handful of candidates, including *DUSP6* which plays context-dependent oncogenic roles such as in glioblastoma, thyroid, and breast cancers, or tumor-suppressive roles in pancreatic, lung, and ovarian cancers(13–15). Our initial observation of aberrantly increased expression of *DUSP6* specifically in sAML HSPCs suggested that DUSP6 may act as an oncogenic driver of leukemic progression. This was corroborated by functional studies including *DUSP6* knockdown experiments which led to reduced viability of *JAK2*-mutant cells. The molecular mechanisms responsible for *DUSP6* induction in the setting of MPN disease evolution are likely to be related to multiple factors, including underlying driver mutations and re-wiring of intracellular signaling pathways in the setting of JAK2 inhibition. Here, we provide proof-of-principle validation of regulatory transcriptional machinery co-mobilized with *DUSP6* in *KLF2* and *KLF1*. Further functional investigations of DUSP6 regulation may be warranted on other targets identified in this study such as EGR family proteins. Notably, we identified increased *DUSP6* expression in *de novo* AML, suggesting that DUSP6 may be a converging downstream effector common to myeloid malignancies. Consistent with this notion, aberrant *FLT3* expression was identified in our initial sAML transcriptome analysis, and FLT3-ITD (a common genetic alteration in *de novo* AML) has been shown to upregulate *DUSP6* expression(61). These findings suggest that DUSP6 may represent a common, targetable dependency in AML.

BCI has been described to be an allosteric inhibitor of both DUSP1 and DUSP6(62) and by this canonical function, elevates ERK signaling. This was observed *in vitro* at low concentrations, but higher BCI doses abrogated pERK, and was concomitant with increased apoptosis and cell cycle arrest. Further pSTAT and pS6 signaling inhibition upon BCI treatment suggest that dual inhibition of these effectors may contribute to the observed functional effects. Similar pathway suppression was also validated upon *DUSP6* knockdown. These observations suggest that rather than off-target effects of BCI, DUSP6 plays novel functions beyond direct regulation of ERK signaling which may be crucial to MPN/AML cell survival. BCI being able to suppress both DUSP1 and DUSP6 have also led to investigations of promiscuity. A recent study suggested a role for DUSP1 in protecting against DNA damage in PV progenitors via modulation of p38/JNK signaling(63). While we cannot exclude the possibility that our findings with BCI may be at least partially related to DUSP1 inhibition, our collective findings including bulk and single cell gene expression analyses, knockdown experiments, and mass cytometry profiling support a predominant role for DUSP6 regulation of MAPK/STAT/S6 phenotypes in leukemic transformation.

The efficacy of BCI has been briefly explored in other hematological malignancies. DUSP1/6 inhibition in combination with FOS inhibitor curcumin and imatinib suppressed tumor growth in BCR-ABL positive chronic myeloid leukemia (CML)(64). Notably in the context of CML, ectopic expression of *DUSP1* but not *DUSP6* conferred resistance to BCI, and inhibitory proliferative effects were a consequence of DUSP1 inhibition, while DUSP6 inhibition favored CML cell survival upon tyrosine kinase inhibitor treatment. Dual pathway inhibition with BCI analog BCI-215 in combination with ruxolitinib also demonstrated durable response in B-cell leukemia (B-ALL) models, where BCI-215 substantially increased ERK phosphorylation, while combination treatment reduced pSTAT5 levels(65). Taken together with the findings from our study, these observations highlight the promising therapeutic potential of DUSP targeting, while reinforcing the notion that DUSP6 plays context-dependent roles even within specific hematologic malignancies. In these settings, the transduction interactome and precise signaling inhibition mechanisms of DUSP6 perturbation remain to be fully characterized.

We also identify *RPS6KA1* as a clinically important marker in AML prognosis and key downstream target of DUSP6. We demonstrate that inhibition of RSK1 via BI-D1870 suppresses S6 signaling *in vitro* and *ex vivo* and exhibits combinatorial efficacy with BCI. BI-D1870 has recently been demonstrated to suppress AML proliferation *in vitro*(66), and a novel RSK inhibitor PMD-026 is currently being evaluated in a phase 1 clinical trial in breast cancer (NCT04115306) in addition to preclinical studies across other cancer types. Thus, our work provides a promising foundation for this novel therapeutic strategy targeting the DUSP6-RSK1-S6 axis in MPNs and AML.

## Acknowledgements

This work was supported by NIH grants R01HL134952 (S.T.O.), and T32HL007088 (J.S.F.). Technical support was provided by the Alvin J. Siteman Cancer Center Tissue Procurement Core Facility, Biostatistics Shared Resource, Flow Cytometry Core, and Immunomonitoring Laboratory, which are supported by NCI Cancer Center Support Grant P30CA91842. The Immunomonitoring Laboratory is also supported by the Andrew M and Jane M Bursky Center for Human Immunology and Immunotherapy Programs. We thank D. Bender, R. Lin, and K. Link for assistance with mass cytometry experiments. We are grateful to Ann Mullally for providing *Jak2* V617F knock-in mice, and to Ross Levine for providing the *MPL* W515L retroviral construct. We thank Mary Fulbright for assistance with mouse colony management. We thank Tim Ley for sharing TCGA LAML data. We thank Feng Gao for assistance with biostatistical analysis. We thank Andreas Vogt for helpful discussions related to BCI. We thank the Genetic Perturbation Service of Goodman Cancer Research Centre at McGill University for access to and preparation of functional genetic tools. G.A.C. was supported by NIH grant R01HL147978, the Edward P. Evans Foundation and the Gabrielle’s Angel Foundation. G.A.C. is a scholar of the Leukemia and Lymphoma Society. This work was also supported by Canadian Institutes of Health Research (CIHR) grants PJT-156233 (S.H.) and PJT-438303 (S.H.). K.Y. is supported by a Canderel Rising Star Summer Studentship and S.H. is supported by a Canadian Research Chair in Functional Genomics.

## Author contributions

T.K., A.B.B.L., K.Y., L.Y., D.A.F, and J.S.F. performed experiments. A.Z W., M.J.A., H.C., G.A.C., and S.T.O. provided technical and clinical support. M.R. evaluated histopathology. T.K., S.H., and S.T.O. designed and supervised the experiments. T.K. and S.T.O. wrote the manuscript. All authors read and approved of the manuscript.

## Competing interests

S.T.O has served as a consultant for Kartos Therapeutics, CTI BioPharma, Celgene/Bristol Myers Squibb, Disc Medicine, Blueprint Medicines, PharmaEssentia, Constellation, Geron, Abbvie, Sierra Oncology, and Incyte. All other authors disclose no competing interests.

## Supplementary figure legends

**Supplementary Figure 1. Elevated DUSP6 expression in patients with sAML compared to MPN.**

A) Heatmap of top differentially expressed genes from microarray analysis in lin- CD34+ cells from sAML patients (n = 14) compared to MF patients (n = 6), and reference expression in healthy donor bone marrow (NBM; n = 5).

B) *DUSP6* expression from CD34+ cells from NBM (n = 31), AML bone marrow CD34+ subfraction (n = 46), and AML bone marrow CD34- subfraction (n = 44) from GSE30029. *DUSP6* values represent quartile normalized, log-transformed values. Statistics were assessed by two-tailed Student’s t test.

C) Immunofluorescence of bone marrow from additional MF and sAML patients, and healthy donors. White arrows denote DUSP6-positive cell staining. Scale bar: 50 µM.

D) Imaging mass cytometry analysis of PBMC cell pellets from normal donor peripheral blood (LRS2), MF (MF20), or sAML (sAML1) patients. Individual images show overlap of indicated channels as denoted. Scale bar = 16 µM.

**Supplementary Figure 2. Serial patient CD34+ scRNA sequencing shows elevated *DUSP6* along MPN to sAML progression.**

A) Gene set enrichment analysis of top altered Hallmark pathways in serial CD34+ samples at the sAML stage compared to MPN stage of three patients.

B) Violin plots showing relative expression of top shared differentiate expressed genes in sAML vs MPN disease states from three patients, and healthy donors (N34, N39).

**Supplementary Figure 3. Correlation between *DUSP6* and top candidates identified along disease progression.**

A) Venn diagram showing shared candidates identified in the top 1000 differentially expressed genes from two sAML stages relative to their chronic MF stage(s).

B) Violin plots highlighting gene expression of shared transcription factors at sAML vs MF stage from from (a).

C) ChIP tracks of key transcription factors identified from scRNA-seq showing occupancy at the *Mus musculus Dusp6* locus across various tissue samples.

D) Heatmap of pearson correlations between *DUSP6* and top, shared candidates across five databases. \**P* < 0.05; \*\**P* < 0.01; \*\*\**P* < 0.001; \*\*\*\**P* < 0.0001.

E) *KLF2* identified as the top correlating gene, and *KLF1* identified among the bottom 10 correlating genes, with *DUSP6* in patient 381812 (MF and sAML) scRNA-seq.

**Supplementary Figure 4. Identification of distinct subpopulations from 381812 CD34+ scRNA-seq.**

A) TotalSeq surface protein detection and mRNA features to guide distinct subpopulation identification.

B) Schematic and relative quantification of distinct subpopulations identified from 381812.

**Supplementary Figure 5. scRNA-seq subpopulation and trajectory analysis of additional patients.**

UMAP analysis and violin plot of *DUSP6*, *KLF2*, and *KLF1* of subpopulations from N34 (and N39) 374024, and 145790. Additional trajectory analysis of patients 374024 and 145790 along disease progression from MPN to sAML.

**Supplementary Figure 6. Exploration of *DUSP6* regulation by *KLF1* and *KLF2* across additional models.**

A) Schematic of a separate serial CD34+ scRNA-seq dataset of a primary MF patient at multiple timepoints along transformation to sAML. As per source publication by Parental *et al.*, sample T1 (PMF) was collected at chronic MPN phase, after which the patient was treated with ruxolitinib for 8 months at which sample T2 (treatment PMF) was collected, and then after 11 months of ruxolitinib treatment at sAML diagnosis (T3; sAML).

B) Violin plots of *DUSP6*, *KLF2*, and *KLF1* expression at different disease timepoints from Parenti *et al.* CD34+ scRNA-seq in (a).

C) Relative fold change of *DUSP6*, *KLF2*, and *KLF1* expression at sAML timepoint (T3) compared to PMF (T1) in across identified cell populations from Parenti *et al.* CD34+ scRNA-seq in (a).

D) *KLF2* (left) and *KLF1* (right) expression from CD34+ cells from NBM (n = 5), MF (n = 14), and sAML (n = 6) patient samples. DUSP6 values represent RMA from microarray. Statistics were assessed by two-tailed Student’s t test.

E) qRT-PCR of *DUSP6* after *KLF2* knockdown in HEL cells. *DUSP6* mRNA expression normalized to *ACTB* for each group and then normalized to pLKO vector control. n = 3 in each group.

F) Immunoblot of DUSP6 expression after *KLF2* knockdown in HEL cells utilizing shRNA or pLKO control vector.

G) Cell viability assay of HEL cells after *KLF2* knockdown relative to control vector. Cells were grown for 96 hours and viability was normalized to the pLKO control vector. n = 6 replicates per construct. Mean and standard deviation presented. Statistics were assessed by two-tailed Student’s t test,

H) *DUSP6* expression in *KLF1* inducible iPSC-derived macrophages relative to control. Dataset investigated: GSE125150.

I) *Klf2* expression in myeloid-specific *Klf2* knockout murine bone marrow derived macrophages. Dataset investigated: GSE149119.

J) Multi-Chip Significance score (S-score) of *Dusp6* in *Klf2* knockout murine embryonic yolk sac erythroid cells. Dataset investigated: GSE27602.

**Supplementary Figure 7. Functional characterization of DUSP6 in MPN/AML.**

A) Relative mRNA expression of DUSP family genes across 35 AML cell lines from the Cancer Cell Line Encyclopedia.

B) Relative mRNA expression of DUSP family genes in HEL cells from the Cancer Cell Line Encyclopedia.

C) Cell viability curves of HEL cells treated with BCI or trametinib across multiple drug doses. Cells were treated for 72 hours and viability was normalized to control. n = 6 replicates at each drug dose. Mean and standard deviation presented.

D) Immunoblot of HEL cells treated with increasing doses of BCI or the MEK inhibitor trametinib. Cells were treated at their indicated drug dose for 24 hours.

E) Phospho-STAT3 and phospho-STAT5 flow cytometry of HEL cells treated with 1µM of BCI or control for 24 hours.

F) Immunoblot profiling of different signaling pathways altered by BCI and trametinib. HEL cells were treated with 1 µM or 1 µM trametinib for 24 hours.

G) Hallmark gene set enrichment analysis showing top altered pathways by normalized enrichment score (NES) and enrichment plots of E2F targets and G2M checkpoint from RNA-seq of HEL cells treated with 1µM of BCI, or DMSO control for 24 hours.

H) Immunoblot of HEL cells ectopically expressing *DUSP6* or *GFP* control vector.

I) Cell viability assay of HEL cells ectopically expressing *DUSP6* or *GFP* control vector. Cells were plated at n= 6 for each condition and grown for 96 hours with viability normalized to the control vector. Mean and standard deviation presented. Statistics were assessed by two-tailed Student’s t test.

J) Cell viability assay of HEL cells ectopically expressing *DUSP6* or *GFP* control vector treated with 300nM BCI. Cells were plated at n= 6 for each condition and grown for 96 hours with viability from normalized to control treatment from each group. Mean and standard deviation presented. Statistics were assessed by two-tailed Student’s t test,

K) Representative images of lin- CD34+ colonies grown in MethoCult H4034 Optimum for 12 days in 0.5 µM BCI or RPMI control. Samples plated in duplicate. Scale bar: 1000 µM.

**Supplementary Figure 8. Suppression of signaling and cytokine production in primary samples by BCI assessed by mass cytometry.**

A) tSNE dimensional reduction clustering of distinct subpopulations from sAML4 and altered signaling upon BCI, TPO induction, or combination treatment post mass cytometry analysis. Samples were treated with 1 µM BCI for 4 hours, 20ng/mL TPO for 1 hour, or combination.

B) TPO-induced signaling across different subpopulations from sAML4. Patient samples were treated with 20ng/mL TPO for 1 hour. Signals were normalized to the control treatment and reported as median Arcsinh ratio.

C) TPO-induced cytokine production across different subpopulations from sAML5. Patient samples were treated with 20ng/mL TPO for 4 hour. Signals were normalized to the control treatment and reported as 90 percentile Arcsinh ratio.

D) Heatmap and dot plots of altered cytokine production of CD14+ monocytes from bone marrow (NBM40) and peripheral blood (NPB LRS2) of healthy donors, and PBMCs from MF and sAML patients by mass cytometry. Unique patient samples were treated with 1 µM BCI for 4 hours, 20ng/mL TPO for 4 hour, or combination. Signals were normalized to the control treatment of each individual patient sample and reported as 90^th^ percentile Arcsinh ratio. Basal cytokine expression in CD14+ cells from MF and sAML are also presented (left panel) and are normalized to the NBM/NPB within each individual CyTOF run to control for batch effect: run 1 - sAML4 and sAML6 normalized to NBM40; run 2 - MF20 and MF102 normalized to NPB LRS2; run 3-MF40 and sAML5 normalized to NBM40.

E) Dot plot of MIP-1β/CCL4 in CD123+ and CD16+ monocyte populations from sAML5. Samples were treated with 1 µM BCI for 4 hours, 20ng/mL TPO for 4 hours, or combination. Signals were normalized to the control treatment and reported as 90 percentile Arcsinh ratio.

**Supplementary Figure 9. Functional characterization of RSK1 in MPN/AML.**

A) Heatmap of inhibitors of upstream regulators of S6 activity and the pearson correlation of their area under curve (AUC) with *DUSP6* expression in AML cell lines.

B) CRISPR dropout screen showing *RPS6KA1* as an essential gene in AML. Candidates were identified if meeting criteria of FDR < 10% and whose inhibition affected # of AML lines but neither of non-AML lines. Data retrieved from Tzelepis *et al*.

C) *RPS6KA1* expression across 10,071 patient samples representing 31 distinct cancer subtypes from the TCGA Pan-Cancer cohort. Expression values provided as log2 (value +1). See additional information in methods.

D) Western blot analysis of *RPS6*KA1 knockdown by shRNA or control vector in HEL cells.

E) Cell viability assay of HEL cells after *RPS6KA1* knockdown relative to control vector. Cells were grown for 96 hours and viability was normalized to the pLKO control vector. n = 6 replicates per construct. Mean and standard deviation presented. Statistics were assessed by two-tailed Student’s t test,

F) Heatmap of altered signaling pathways of lin- CD34+ cells from unique normal bone marrow donors and peripheral blood of MF patients by mass cytometry. Patient samples were treated with 5 µM BI-D1870 for 4 hours, 20ng/mL TPO for 1 hour, or combination. Signals were normalized to the control treatment of each individual patient sample and reported as 90^th^ percentile Arcsinh ratio.

G) Ridge plot of *RPS6KA1* expression from CD34+ scRNA-seq of N34, N39, and 381812 at MF and sAML stages.

H) *RPS6KA1* expression from CD34+ cells from NBM (n=5), MF (n=14), and sAML (n=6) patient samples. *RPS6KA1* values represent RMA from microarray. Statistics were assessed by two-tailed Student’s t test.

I) Cell viability curve of HEL cells after ectopic expression of *RPS6KA1* or *GFP* control treated with increasing concentrations of BI-D1870. Cells were treated for 96 hours and viability was normalized to the control treatment from each group. n = 6 replicates per construct. Mean and standard deviation presented.

J) Cell viability assay of HEL cells treated with 1 µM BI-D1870, 300 µM BCI or combination, and UKE-1 cells treated with 2 µM BI-D1870, 200 µM BCI, or combination. Cells were treated for 72 hours and viability was normalized to the control treatment. n = 6 replicates at each drug dose. Mean and standard deviation presented.

K) Hallmark gene set enrichment analysis showing top altered pathways from RNA-seq of HEL cells treated for 24 hours with 10 µM BI-D1870 + 1 µM BCI compared to DMSO control (left) and 10 µM BI-D1870 + 1 µM BCI compared to 10 µM BI-D1870 alone (right).

L) Dot plot of mass cytometry analysis of lin- CD34+ cells from MF103 treated with 1 µM BCI for 4 hours, 5 µM BI-D1870 for 4 hours, 20ng/mL TPO for 1 hour, or combination. Signals of key phosphorylated proteins were normalized to the control treatment and reported as 90^th^ percentile Arcsinh ratio.

**Supplementary Figure 10. DUSP6 mediates response to JAK2 inhibitors.**

A) Correlation of *DUSP6* expression and fedratinib IC_50_ in 18 AML cell lines. *DUSP6* expression obtained from the CCLE database and fedratinib IC_50_ obtained from the GDSC1 collection.

B) Cell viability assay of HEL parental or HEL Fed-P cells treated with BCI at the indicated doses. Cells were treated for 72 hours and viability was normalized to the control treatment from each group. n = 6 replicates at each drug dose. Mean and standard deviation presented.

C) Cell viability assay of HEL cells treated with BCI, fedratinib, or combination. Cells were treated for 72 hours at the indicated drug doses and viability was normalized to the control treatment. n = 6 replicates at each drug dose. Mean and standard deviation presented.

**Supplementary Figure 11. BCI alleviates disease burden across MPN and sAML mouse models.**

A) WBC subpopulation counts, platelet counts, and body weight from *Jak2* transplanted mice treated with vehicle or BCI across multiple timepoints. Statistics were assessed by two-way ANOVA comparing vehicle to BCI.

B) Representative gross spleen of *Jak2* mice treated with vehicle or BCI at endpoint.

C) Hematocrit, white blood cell (WBC) counts and differentials, and platelets counts of wildtype primary mice treated with vehicle (n = 4) or 25 mg/kg BCI (n = 5) following weekly schedule of 5 days on, 2 days off treatment across multiple timepoints. Liver, spleen, and body weights collected at endpoint. Two-way ANOVA and two-tailed Student’s t test statistical analysis resulted in non-significant values across all comparisons between vehicle and BCI treatment.

D) Representative flow cytometry analysis of peripheral blood from CD45.1 mice showing engraftment of *MPL* W515 GFP+ CD45.2 cells.

E) WBC subpopulation counts, platelet counts, and normalized spleen weight from *MPL* W515 MF model of transplanted mice treated with vehicle or BCI across multiple timepoints. Statistics were assessed by two-way ANOVA for white count differential comparisons between vehicle and BCI and two-tailed Student’s t test for normalized spleen weight.

F) Normalized spleen and liver weights from mice at end-point from sAML14 CD34+ PDX.

G) tSNE dimensional reduction clustering of mouse and human CD45+ cells from bone marrow of sAML14 PDX mice.

---

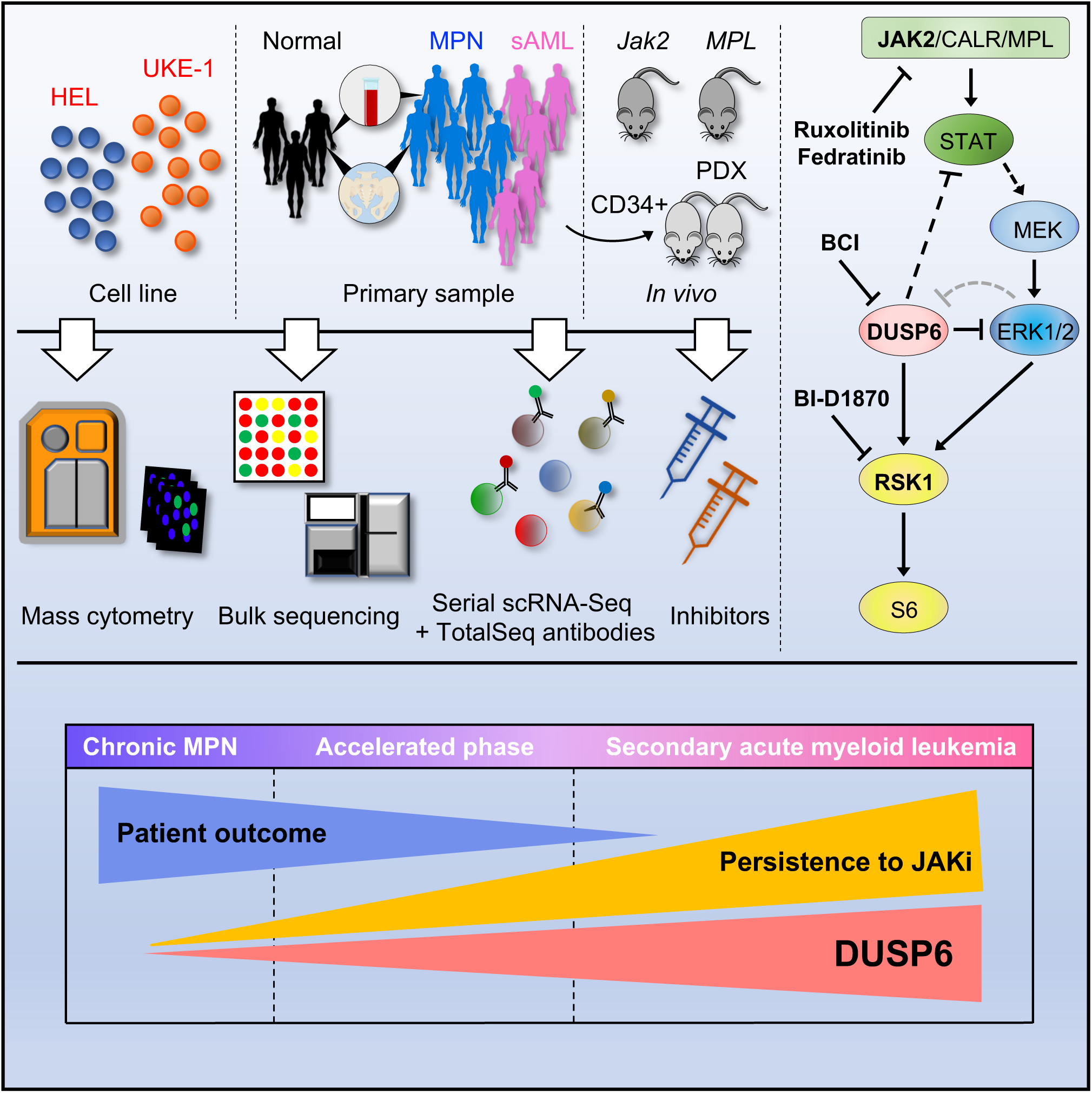

