## Supplementary Figures 1-11 for "DUSP6 mediates resistance to JAK2 inhibition and drives leukemic progression"

### Supplementary Figure 1

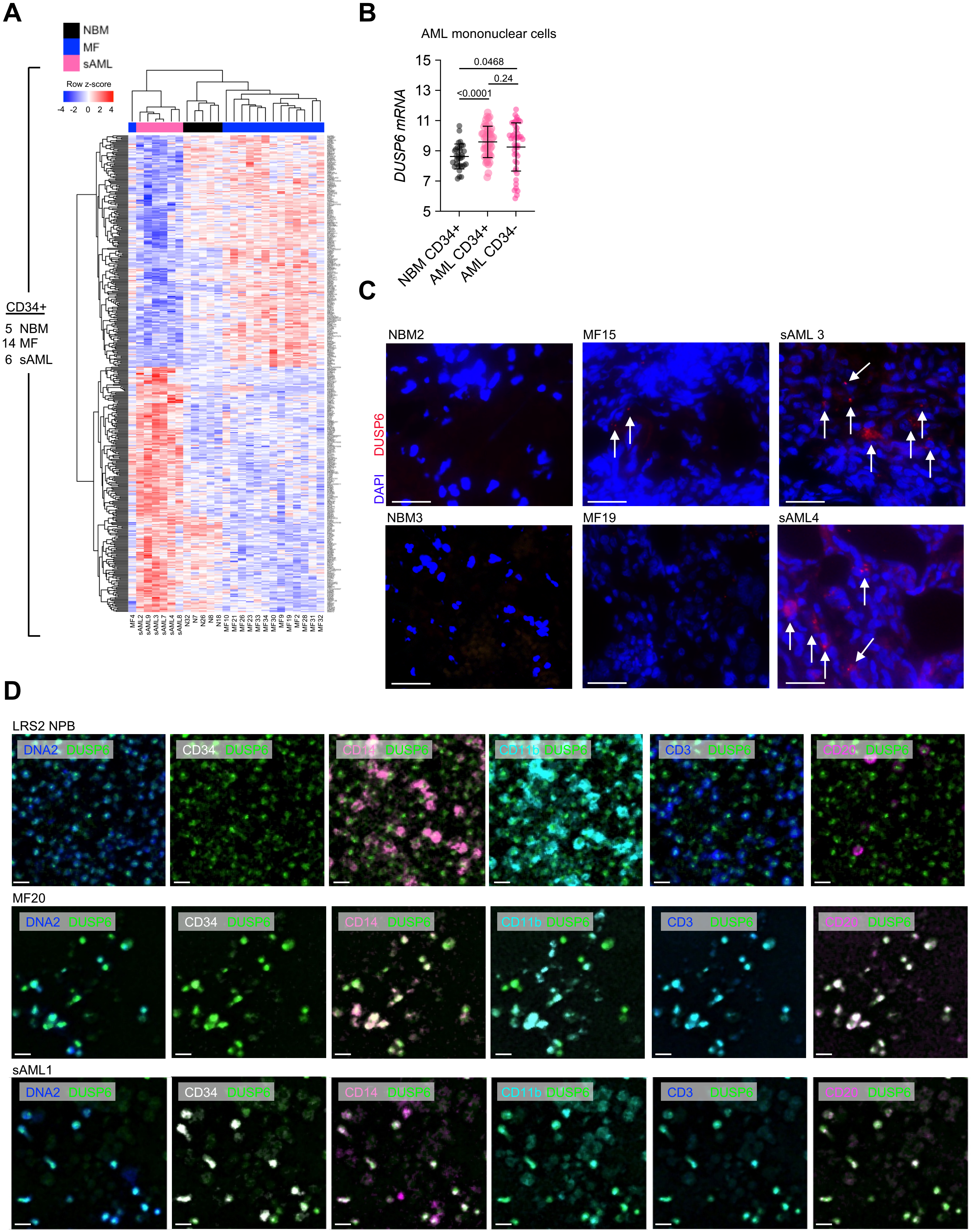

### Supplementary Figure 2

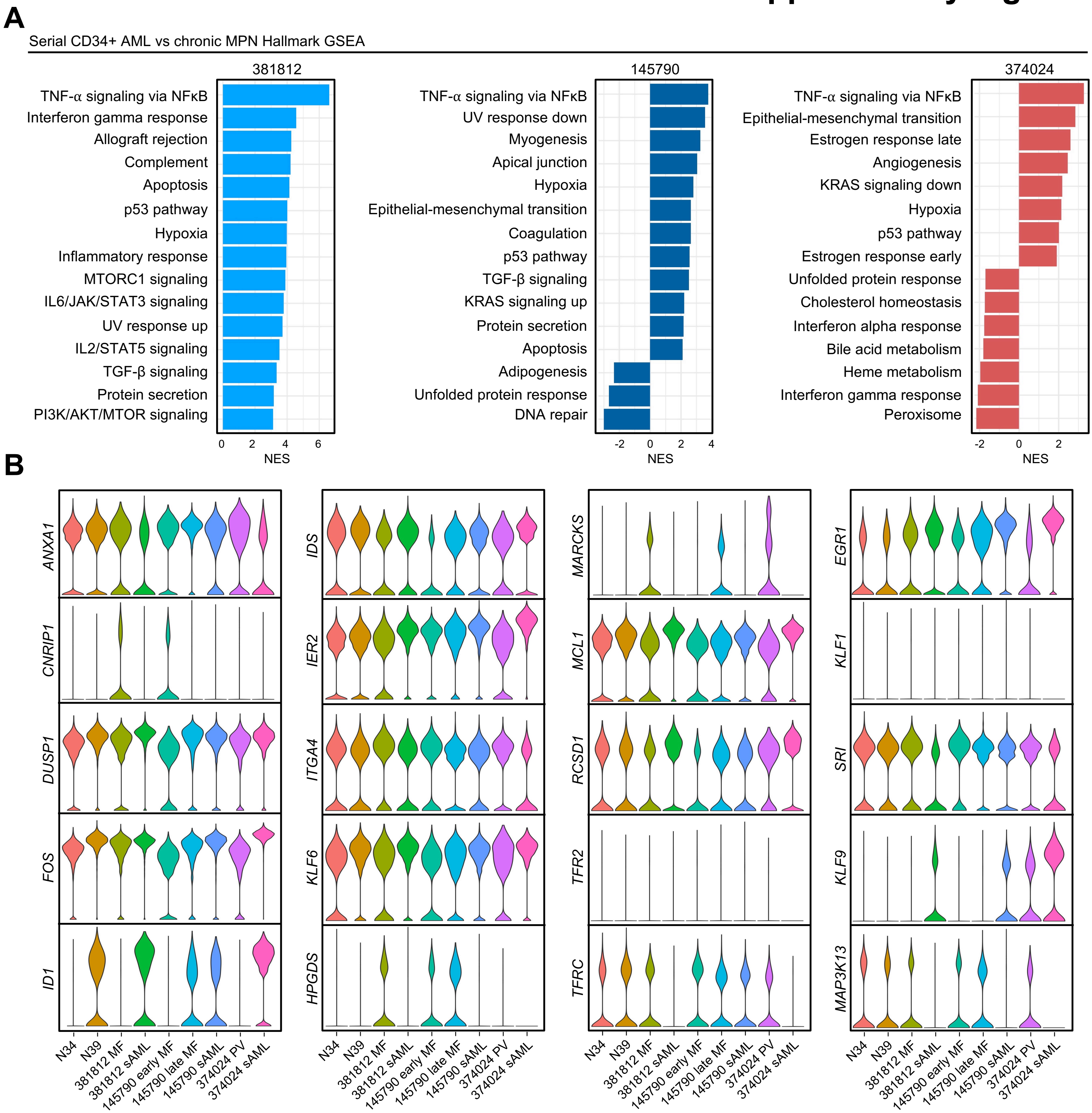

### Supplementary Figure 3

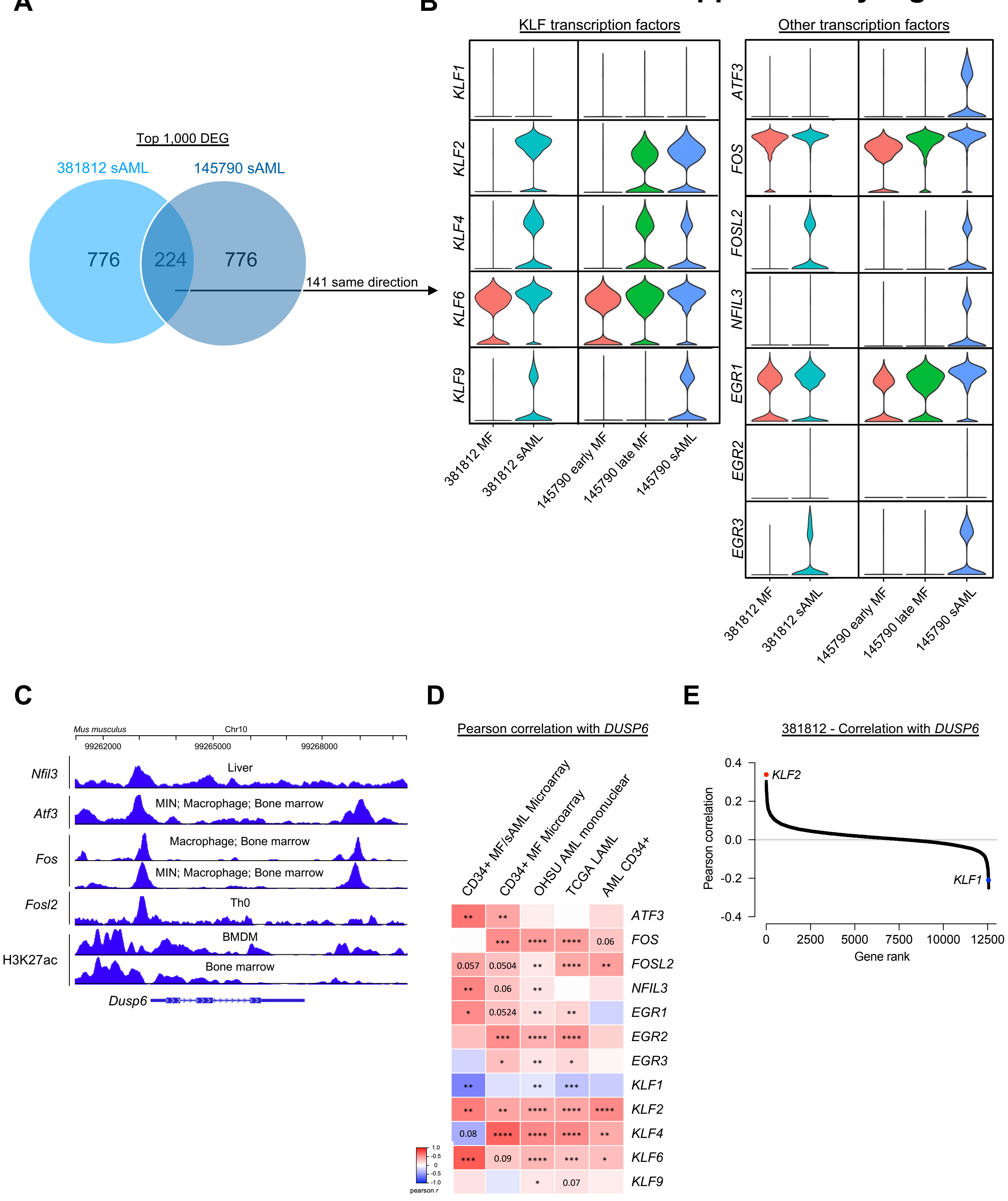

### Supplementary Figure 4

**A**

381812

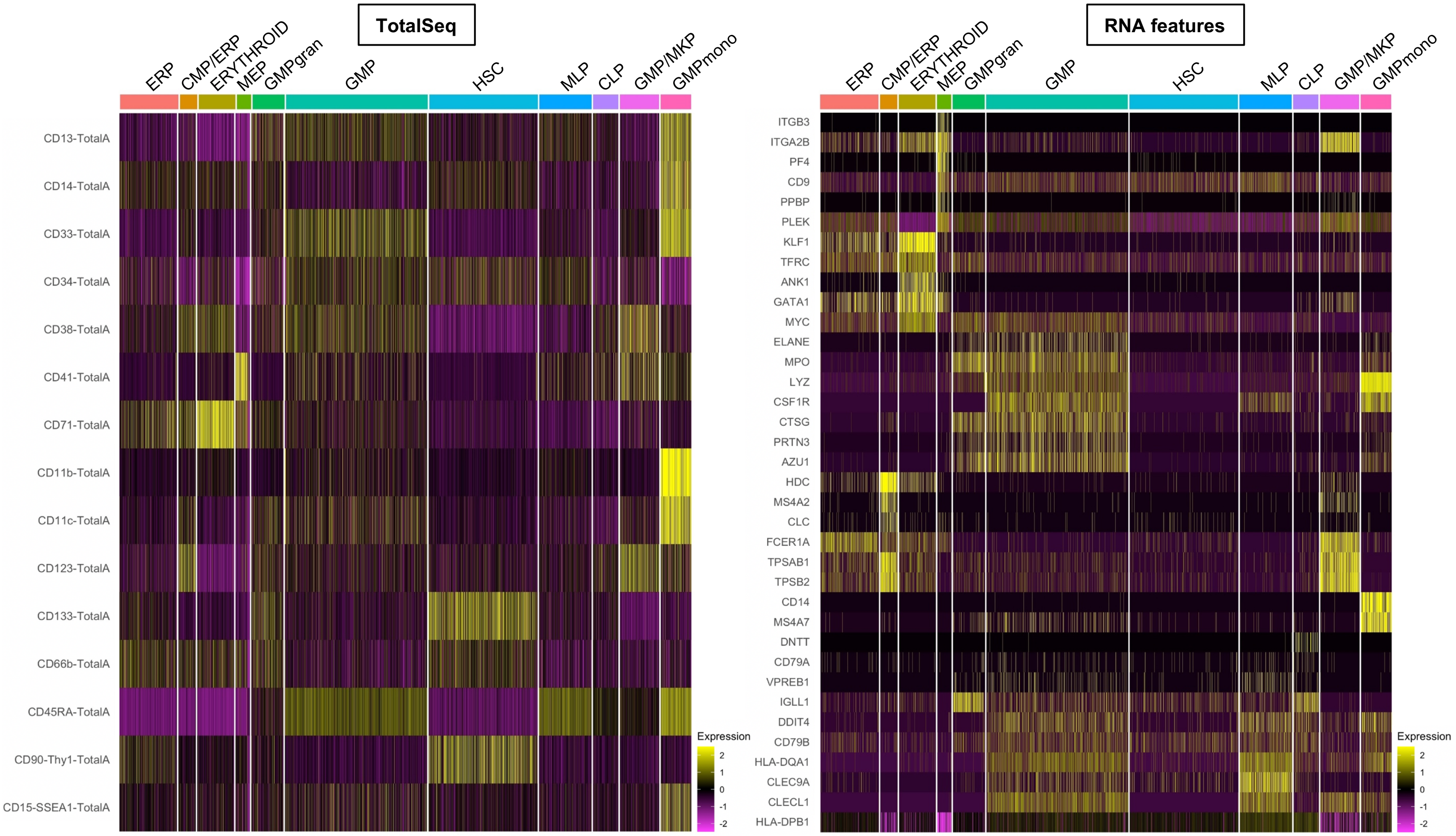


**B**

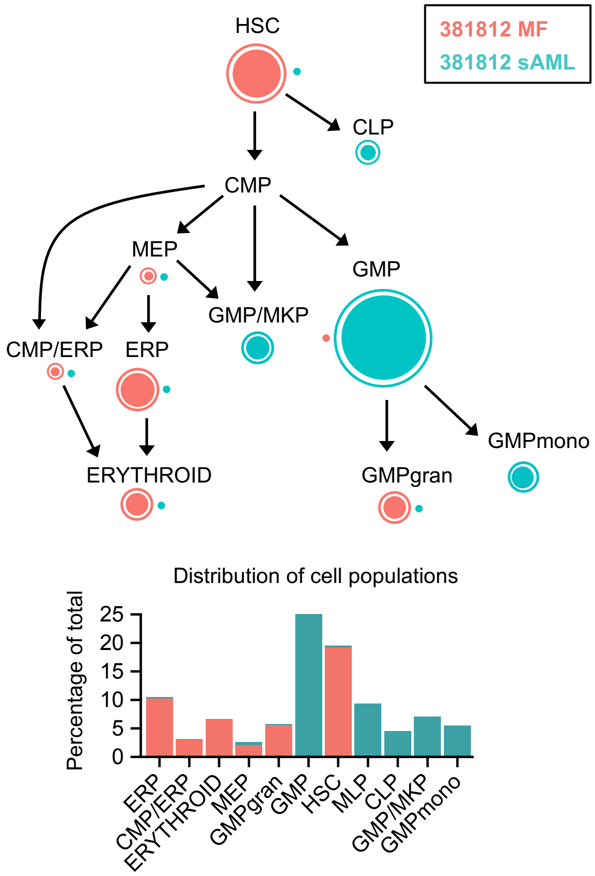

#### Supplementary Figure 5

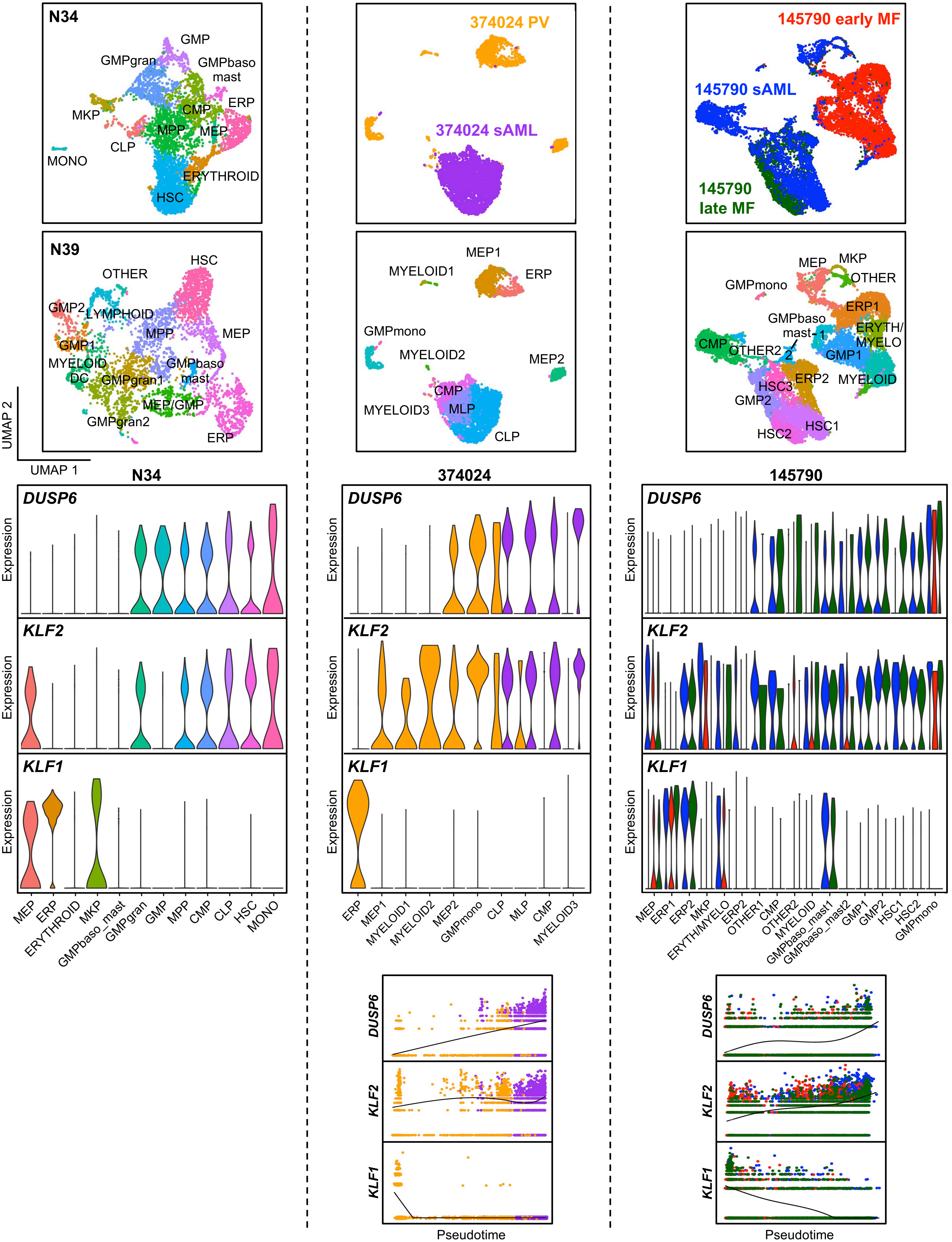

### Supplementary Figure 6

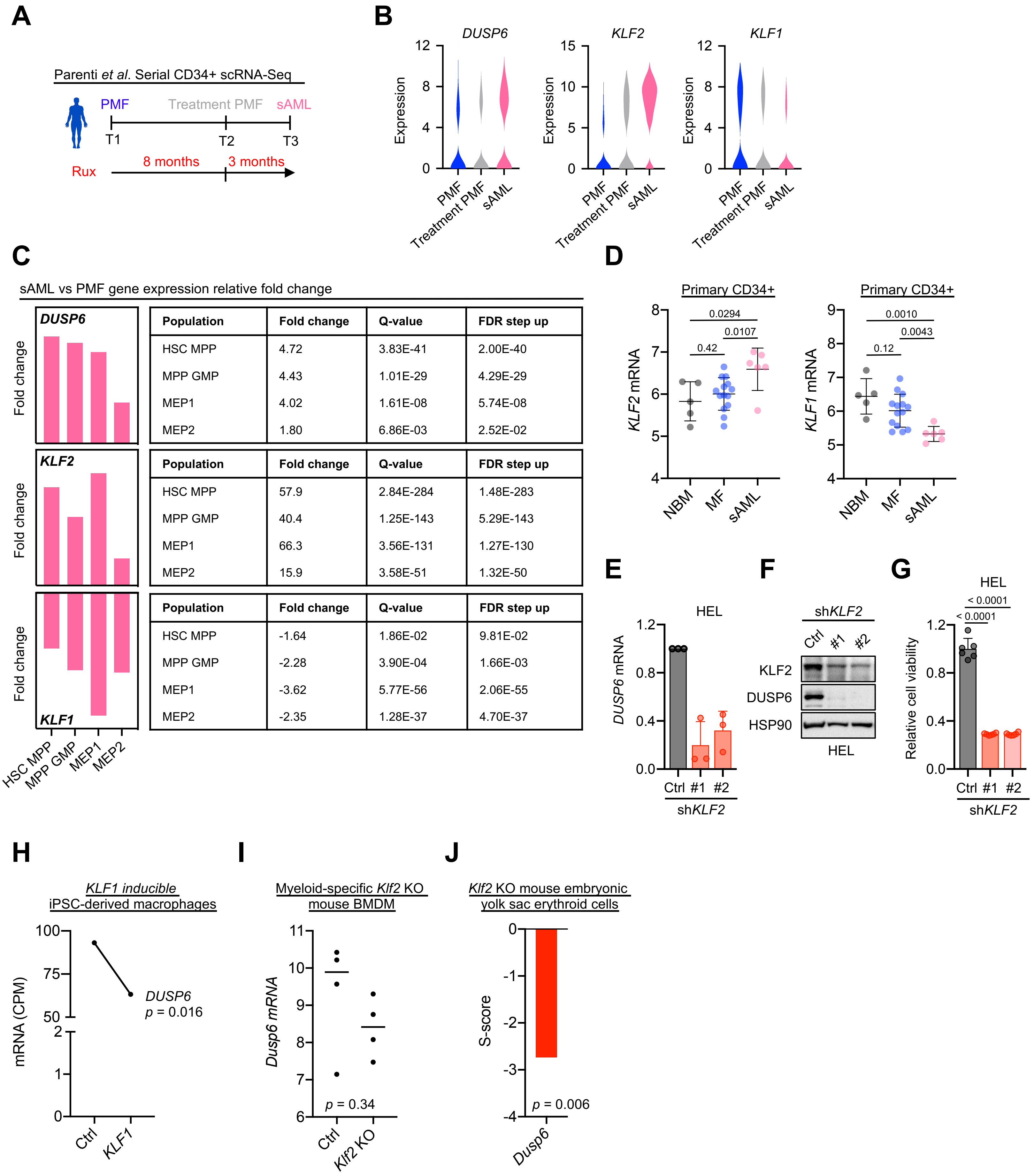


**DUSP6**

| Population | Fold change | Q-value | FDR step up |
|---|---|---|---|
| HSC MPP | 4.72 | 3.83E-41 | 2.00E-40 |
| MPP GMP | 4.43 | 1.01E-29 | 4.29E-29 |
| MEP1 | 4.02 | 1.61E-08 | 5.74E-08 |
| MEP2 | 1.80 | 6.86E-03 | 2.52E-02 |

**KLF2**

| Population | Fold change | Q-value | FDR step up |
|---|---|---|---|
| HSC MPP | 57.9 | 2.84E-284 | 1.48E-283 |
| MPP GMP | 40.4 | 1.25E-143 | 5.29E-143 |
| MEP1 | 66.3 | 3.56E-131 | 1.27E-130 |
| MEP2 | 15.9 | 3.58E-51 | 1.32E-50 |

**KLF1**

| Population | Fold change | Q-value | FDR step up |
|---|---|---|---|
| HSC MPP | -1.64 | 1.86E-02 | 9.81E-02 |
| MPP GMP | -2.28 | 3.90E-04 | 1.66E-03 |
| MEP1 | -3.62 | 5.77E-56 | 2.06E-55 |
| MEP2 | -2.35 | 1.28E-37 | 4.70E-37 |

### Supplementary Fig. 7

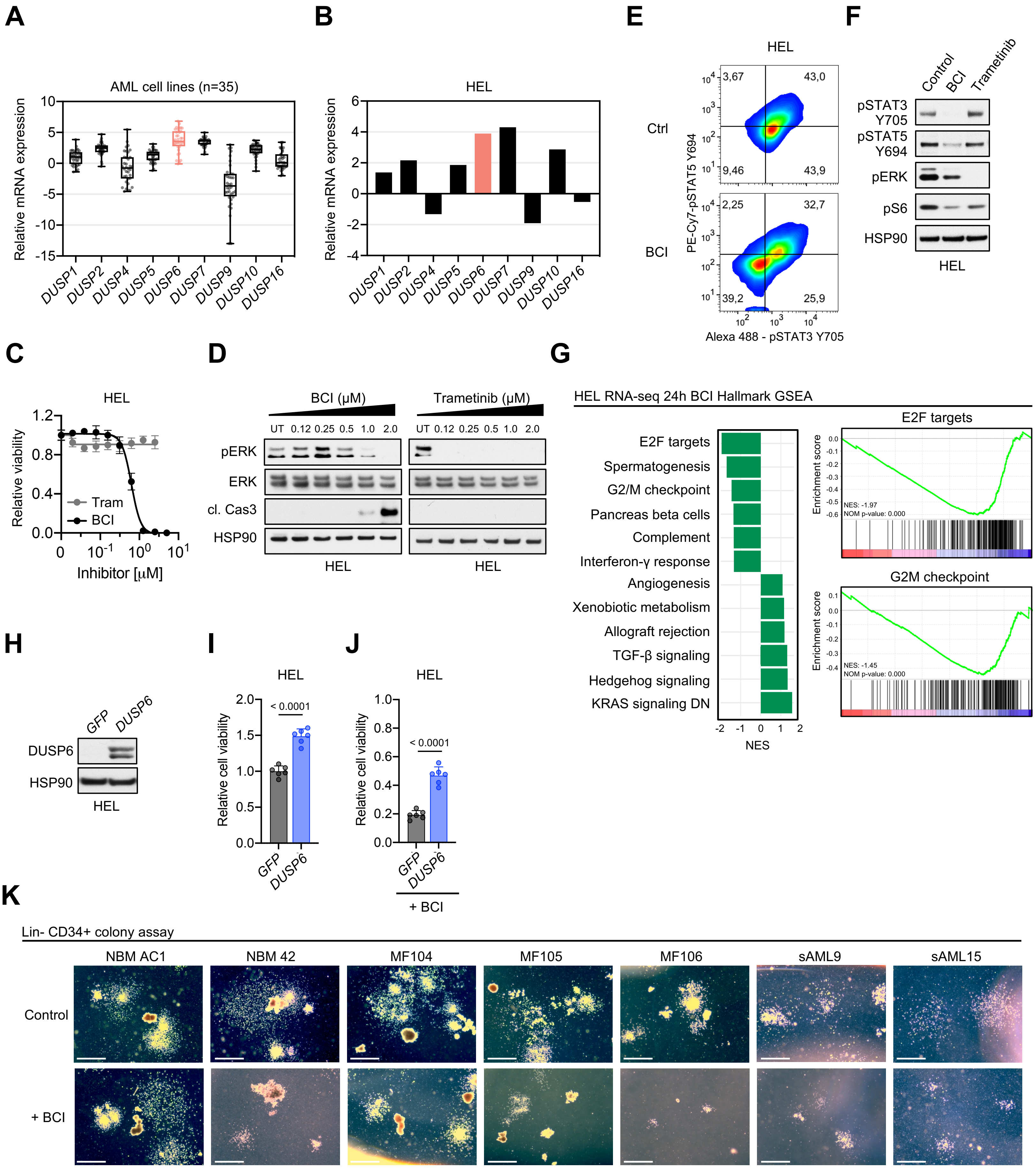

### Supplementary Figure 8

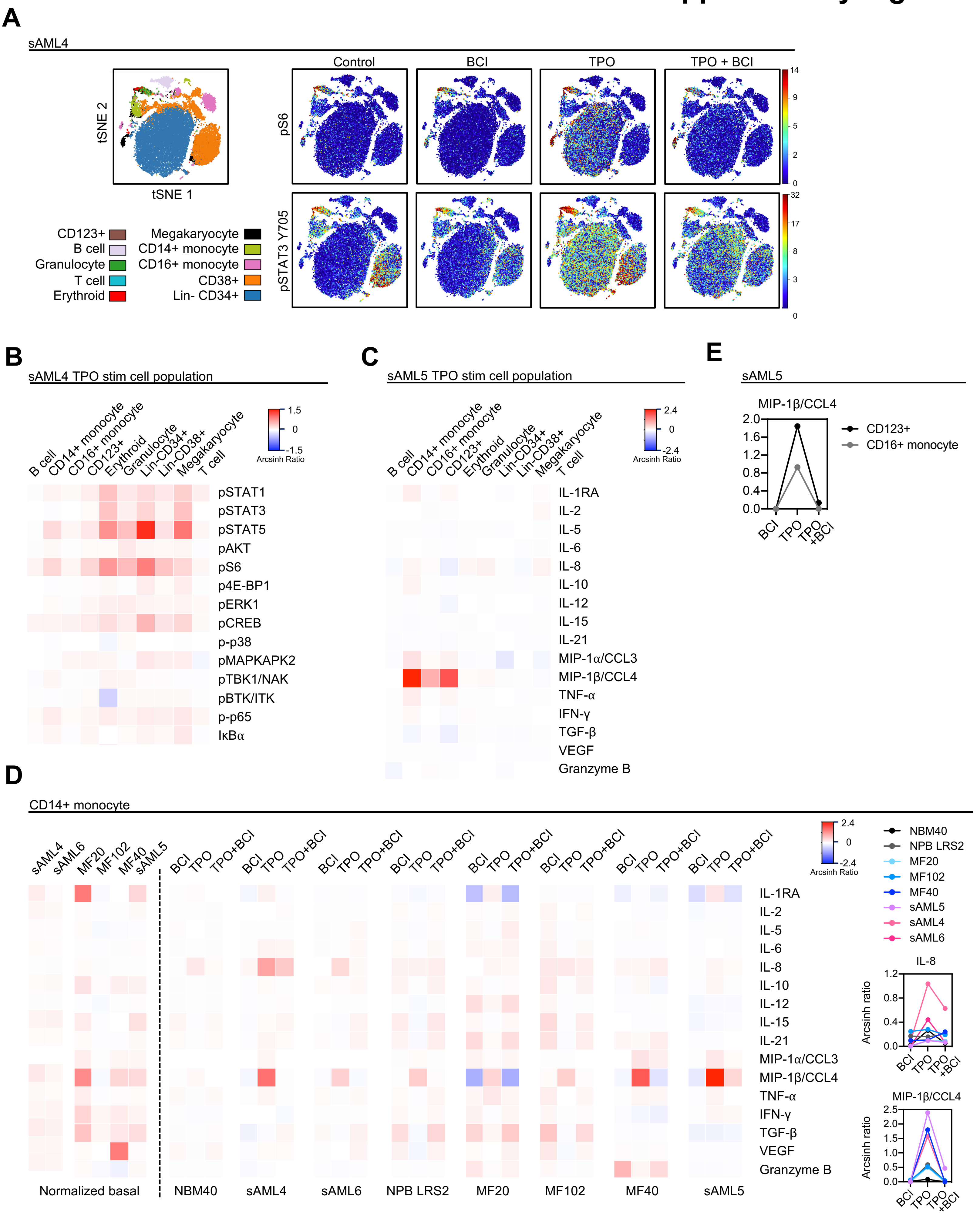

### Supplementary Figure 9

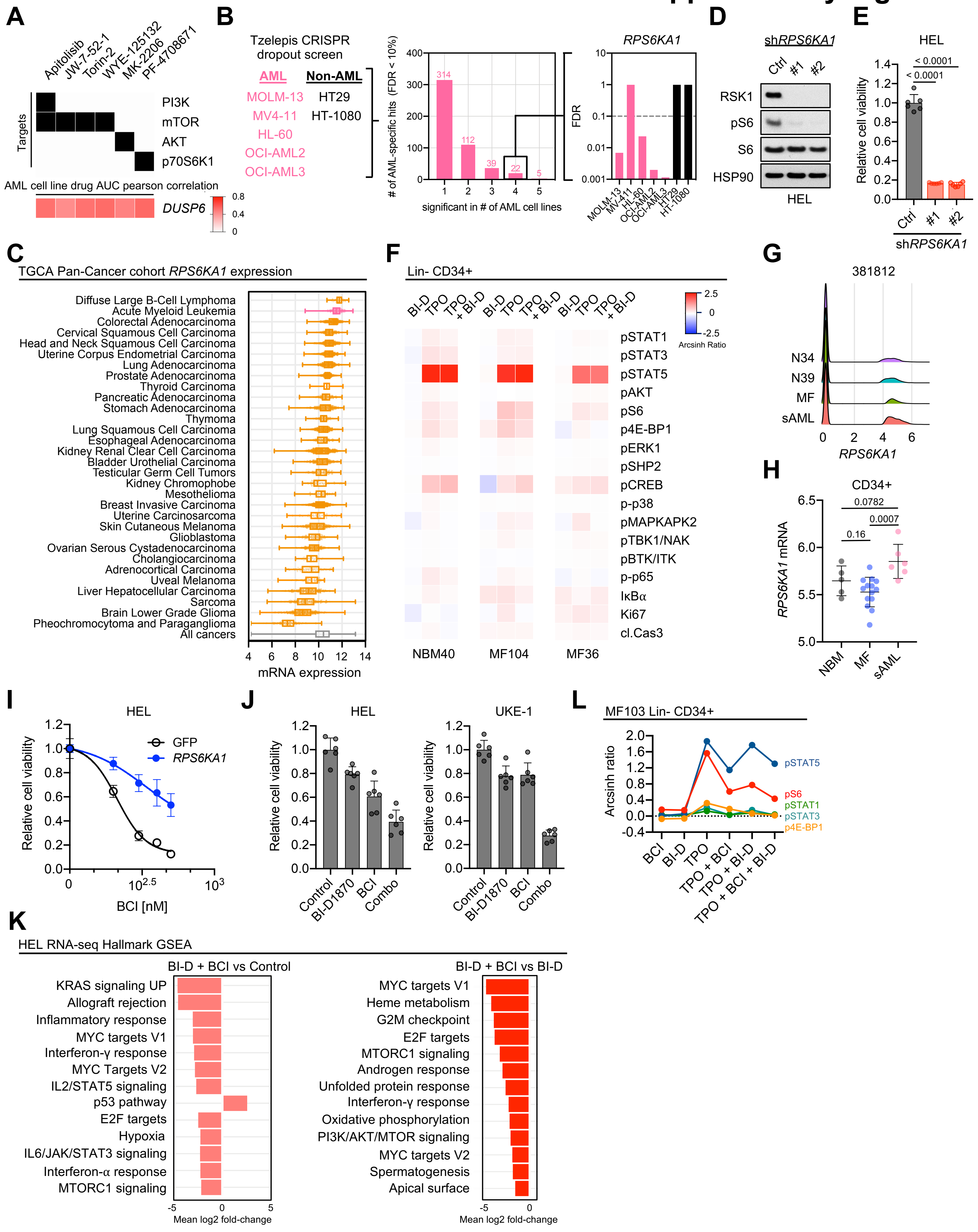

### Supplementary Figure 10

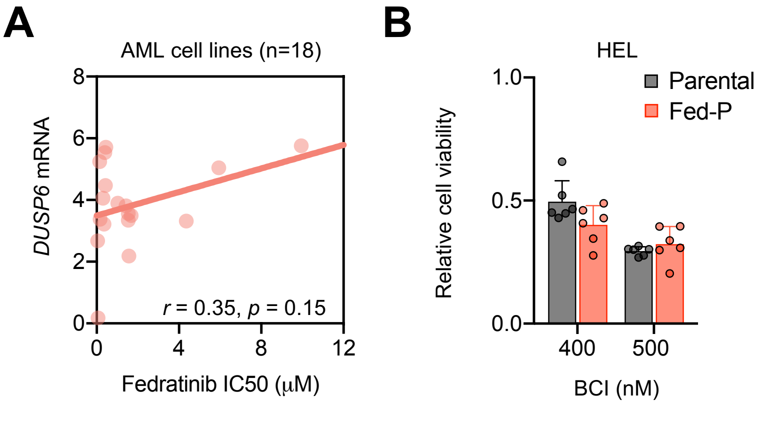

### Supplementary Figure 11

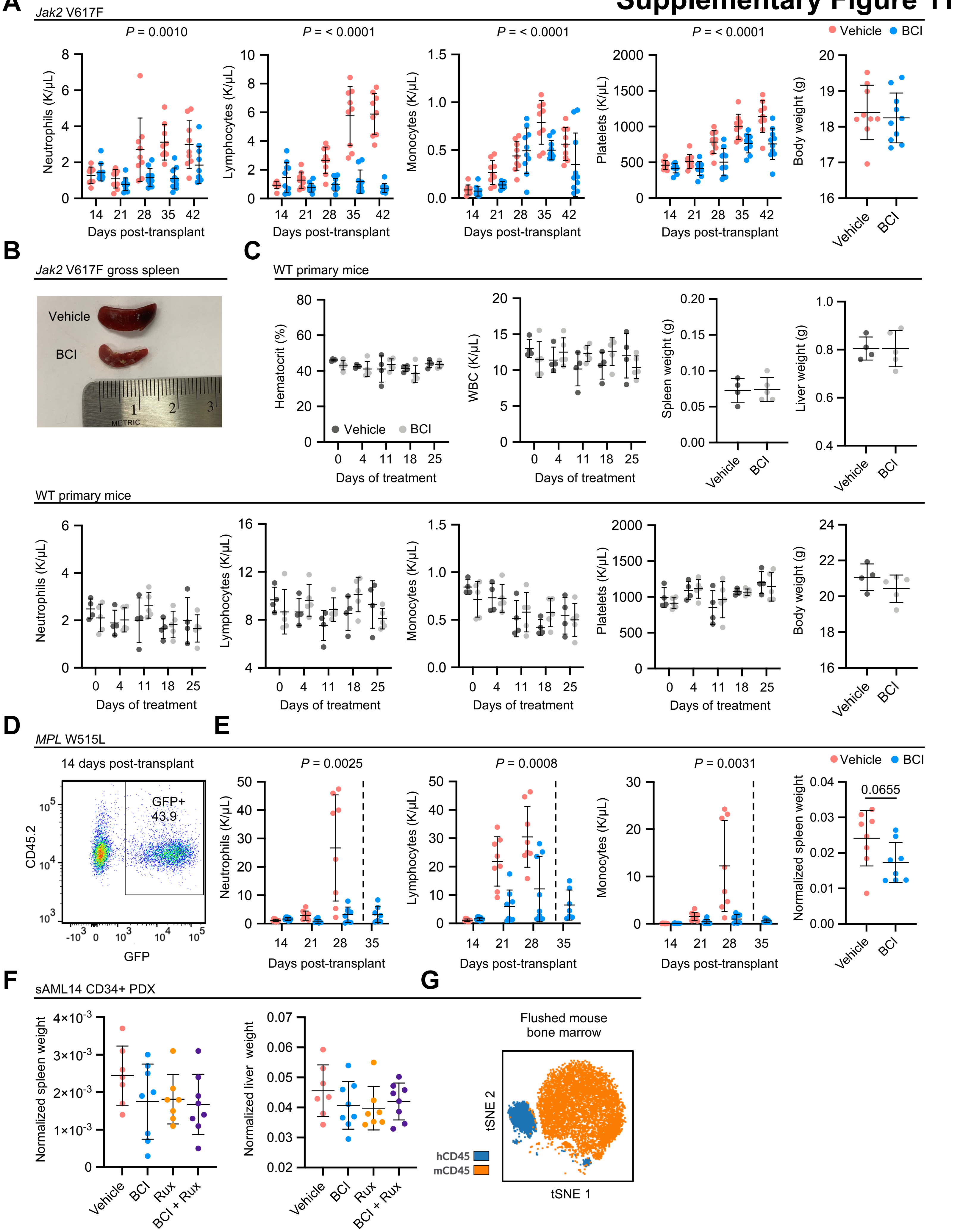
